# Control of clathrin-mediated endocytosis by NIMA family kinases

**DOI:** 10.1101/757427

**Authors:** Braveen B. Joseph, Yu Wang, Phil Edeen, Vladimir Lažetić, Barth D. Grant, David S. Fay

## Abstract

Endocytosis, the process by which cells internalize plasma membrane and associated cargo, is regulated extensively by posttranslational modifications. Previous studies suggested the potential involvement of scores of protein kinases in endocytic control, of which only a few have been validated within their native context. Here we show that the conserved NIMA-related kinases NEKL-2/NEK8/9 and NEKL-3/NEK6/7 (the NEKLs) control clathrin-mediated endocytosis in *C. elegans*. Loss of NEKLs leads to clathrin mislocalization and to a dramatic reduction in clathrin mobility at the apical membrane. Strikingly, reducing the levels of active AP2, the major clathrin adapter complex, rescues *nekl* mutant defects, whereas increased levels of active AP2 exacerbate nekl defects. Moreover, NEKL inhibition alleviates defects associated with reduced AP2 activity, attesting to the tight link between NEKL and AP2 functions. We also show that NEKLs are required for the clustering and internalization of membrane cargo and that human NEKs rescue defects in *nekl* mutant worms.

## Introduction

The cuticle of *C. elegans* is a flexible apical extracellular matrix consisting of cross-linked collagens, non-collagenous proteins, linked carbohydrates, and lipids [1, 2]. The cuticle is essential for providing a protective barrier from the environment, for maintaining the proper shape and integrity of the organism, and for facilitating muscle-based locomotion by functioning as an exoskeleton [3]. Remodeling of the cuticle occurs at the end of each of four larval stages (L1–L4) through a process called molting. During molting, a new cuticle is synthesized under the old cuticle, which is then shed [3-5]. Molting enables organismal growth and allows for changes in the composition and organization of the cuticle at different life stages. Molting defects can occur when synthesis of the new cuticle is compromised or when shedding of the old cuticle is incomplete. A sizeable number of factors have been implicated in *C. elegans* molting including proteins involved in cuticle structure, protein modification, protein degradation, cell signaling, transcription, and intracellular trafficking [4, 6].

During each molt, an accumulation of ribosomes, Golgi bodies and RNA is observed within the epidermal cells that produce the new cuticle, consistent with increased protein synthesis [1, 2, 5, 7]. The secretion of essential structural proteins, along with the enzymatic activities required for cuticle replacement, is accomplished through exocytosis. Consistent with this, inhibition of *sec-23*, which encodes a component of COPII-coated vesicles required for the transport of proteins from the endoplasmic reticulum (ER) to the Golgi, leads to molting defects [8]. At the same time, endocytosis is required to balance exocytosis and thus maintain a relatively constant volume/area of apical plasma membrane. In addition, the recycling of old cuticle components may be enabled through the process of endocytosis.

Endocytosis by the epidermis is also essential for the uptake of sterols from the environment, which provide building blocks for the hormonal cues that drive molting [4, 6, 9-12]. Consistent with this, worms deprived of cholesterol fail to molt [13-15]. Sterol uptake by the epidermis is thought to be dependent in part on LRP-1 (human LRP2), which belongs to the low-density lipoprotein (LDL) receptor family of integral membrane proteins. Inhibition of LRP-1 and other trafficking components required for LRP-1 uptake, such as the adapter protein DAB-1 (human DAB1/2), also lead to defective molting [15-19].

We previously reported that knockdown of NEKL-2 (human NEK8/9) or NEKL-3 (human NEK6/7), two conserved members of the Never-In-Mitosis-A (NIMA) protein kinase family, leads to molting defects in *C. elegans* [20, 21]. In addition, loss of function in the conserved ankyrin repeat proteins MLT-2 (human ANKS6), MLT-3 (human ANKS3), and MLT-4 (human INVS), leads to molting defects that are identical to those of *nekl* mutants. The NEKL–MLTs form two distinct complexes (NEKL-2 with MLT-2–MLT-4 and NEKL-3 with MLT-3), and the MLTs are required for the correct subcellular localization of the NEKLs [21]. Importantly, these physical and functional interactions between NEKLs and MLTs appear to be highly conserved [22-25].

In mammals, the orthologs of NEKL-2 and NEKL-3 have mainly been studied in the context of cell cycle progression (NEK6, NEK7, NEK9) [26-30] and ciliogenesis (NEK8 and NEK9) [25, 31-36] and have correspondingly been implicated in a spectrum of cancers and ciliopathies [23, 31, 34, 37-49]. However, high-throughput screens also identified NEK6, NEK7, NEK8, and NEK9 as candidate regulators of clathrin-mediated endocytosis [50, 51], although these findings have not been validated. Furthermore, in *Aspergillus nidulans* and *Saccharomyces cerevisiae*, NIMA orthologs have been shown to functionally interact with ESCRT genes, which promote membrane remodeling and endosome maturation [52, 53].These results suggest that NEK family members may have conserved roles in intracellular trafficking that have been largely overlooked.

We previously showed that NEKL–MLTs are expressed in punctate patterns in the major syncytial epidermis of the worm, hyp7, suggesting that NEKL–MLTs localize to one or more trafficking compartments [20, 21]. In addition, we found that molting-defective *nekl–mlt* larvae exhibit abnormal morphology and/or localization of multiple trafficking markers in the apical region of hyp7, including a multi-copy clathrin heavy chain reporter [20, 21]. However, it was unclear as to whether the apparent defects in clathrin-mediated endocytosis were a primary cause of the observed molting defects in *nekl–mlt* mutants or a secondary consequence of physiological effects caused by the presence of a double cuticle and the inability of the worms to feed.

Clathrin-mediated endocytosis is a highly regulated stepwise process involving dozens of factors that act temporally to control the initiation, maturation, and internalization of clathrin-coated pits/vesicles [54-60]. Among these are the components of the clathrin scaffold itself, multiple triskelion units containing three heavy chains and an associated light chain. In addition, clathrin-mediated endocytosis requires numerous adapter proteins that link clathrin to the plasma membrane and to integral membrane cargo. Chief among these is the conserved plasma membrane adapter protein complex, AP2 [57, 61-63]. AP2 consists of four subunits, termed α, β, µ, and σ, and exists in at least two functionally distinct structures, broadly termed the open/active and closed/inactive conformations [55, 57, 64-67]. Allosteric regulators of AP2 conformation include FCHo1/2, which promotes the open state of AP2 [68, 69], and NECAP1/2, which promotes the closed conformation [70]. In addition, protein kinases have been implicated in the regulation of AP2 through phosphorylation of the µ subunit [67, 71].

Here we demonstrate that NEKL–MLTs regulate clathrin-mediated endocytosis within the context of an intact developing organism. We show that the function of NEKL–MLTs in trafficking is highly sensitive to the balance between the open and closed AP2 conformations and that AP2-associated phenotypes are also responsive to NEKL activity. In addition, we demonstrate that loss of NEKL functions leads to defects in LRP-1/LRP2 endocytosis, a cargo that is physiologically relevant to molting. Our combined findings indicate that defects in endocytosis are likely to be a major underlying basis for the observed molting defects in *nekl* mutants. Finally, we show that mammalian NEK6 and NEK7 can partially rescue endocytosis and molting defects in *nekl-3* mutant worms.

## Results

### *nekl–mlt* defects are suppressed by decreased function of the AP2 clathrin-adapter complex

We previously described a genetic and bioinformatic approach to identify suppressors of larval lethality in strains deficient for NEKL kinase activity [72]. Our screen makes use of weak aphenotypic alleles of *nekl-2(fd81)* and *nekl-3(gk894345)*, which, when combined in double mutants, lead to penetrant larval arrest due to molting defects [72, 73]. Homozygous *nekl-2(fd81); nekl-3(gk894345)* mutants (hereafter referred to as *nekl-2; nekl-3* mutants) can be propagated only in the presence of a *nekl-2*^*+*^ or *nekl-3*^*+*^ rescuing extrachromosomal array, whereas strains that acquire a suppressor mutation no longer require the array for viability (Fig 1A,B).

**Fig 1.**
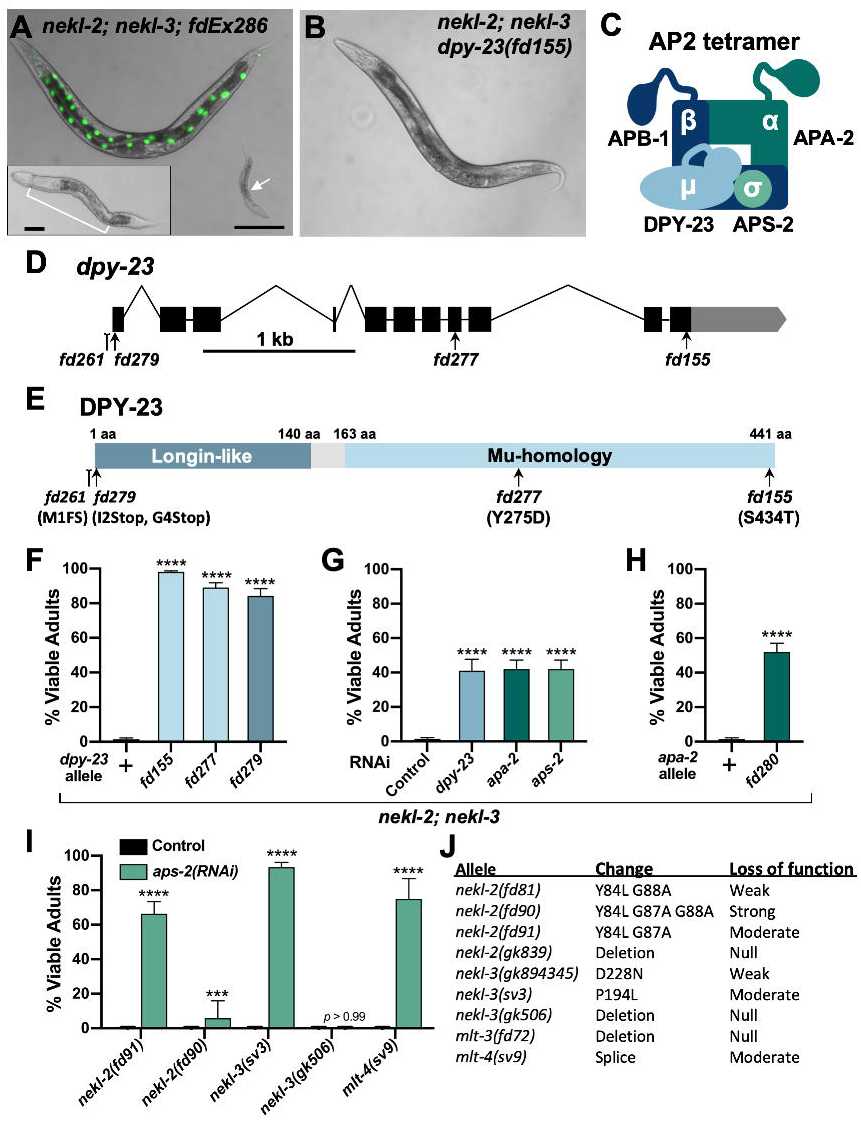
Loss of AP2 complex activity suppresses *nekl–mlt* molting defects. (A) Merged DIC and fluorescence images of *nekl-2(fd81); nekl-3(gk894345)* worms. The rescuing extrachromosomal array (*fdEx286*) expresses wild-type *nekl-3* and the *SUR-5::GFP* reporter. White arrow indicates a growth-arrested *nekl-2*; *nekl-3* larva that failed to inherit the array and exhibits a “corset” morphology, characteristic of *nekl–mlt* molting defects (inset in A). The bracket marks the constricted mid-body region, which contains a double cuticle. Bar size in A = 100 µm (for A and B); in inset, 20 µm. (B) DIC image of *nekl-2*; *nekl-3* adult worm containing the suppressor mutation *dpy-23(fd155)*. (C) Graphic representation of AP2 tetramer complex containing four subunits. (D) Gene structure diagram of *dpy-23* including the locations of mutations. Point mutations *(fd155, fd277, fd279)* are indicated by arrows; indel *(fd261)* by the line ending in a small bracket. (E) Protein domain diagram of DPY-23 with corresponding allelic changes; *fd261* is missing ∼30 bp of the proximal 5’UTR including a predicted SL1 transplice site and the start codon. The amino acid (aa) locations of the two domains are indicated. (F–I) Bar plots showing suppression of molting defects in *nekl–mlt* mutants by reduction in AP2 activity. Assays F–H were carried out in *nekl-2(fd81)*; *nekl-3(gk894345)* double mutants. (G,I) RNAi was carried out in the indicated backgrounds using injection methods; control indicates non-injected siblings. (F–I) Error bars indicate 95% confidence intervals; p-values were determined using Fischer’s exact test where proportions were compared to the wild-type allele (F,H) or to the RNAi control (G,I): \*\*\*\**p* < 0.0001, \*\*\**p* < 0.001. (J) Guide to *nekl–mlt* alleles used in this study. Raw data are available in S1 File.

Among *nekl-2; nekl-3* animals containing the *fd155* suppressor mutation, 98% progressed to adulthood versus only 1–2% for the parental *nekl-2; nekl-3* strain (Fig 1B,F). We showed the *fd155* causal mutation to be a T-to-A transversion in the 11^th^ exon of *dpy-23*, which leads to a S434T substitution in a serine that is highly conserved (Fig 1D,E). A second independent mutation (*fd277*) was identified as a T to G transversion in the 8^th^ exon of *dpy-23*, leading to a Y275D substitution in a highly conserved tyrosine residue (Fig 1D,E). Both mutations, as well as a CRISPR/Cas9-generated predicted null allele of *dpy-23 (fd279*), led to a high percentage of viable adults in the nekl-2; nekl-3 background (Fig 1D–F; S1 Table), and partial suppression was observed in *nekl-2; nekl-3* strains treated with *dpy-23(RNAi*) (Fig 1G).

*dpy-23* (also known as *apm-2*) encodes the µ subunit of the *C. elegans* AP2 complex [74, 75]; the other three subunits are encoded by *apa-2* (α), *apb-1* (β), and *aps-2* (σ) (Fig 1C) [57, 76]. AP2 binds to phosphatidylinositol-4,5-bisphosphate (PIP2) lipids on the plasma membrane and functions as an adapter, linking cytoplasmic clathrin to plasma membrane cargo [57, 76]. In *C. elegans*, AP2 subunits form two partially independent hemicomplexes composed of µ/β and α/σ [77]. Although normal levels of clathrin-mediated endocytosis occur only when all four subunits are functional, *C. elegans* strains containing either the µ/β or α/σ hemicomplex are nevertheless viable, as demonstrated by the ability to propagate homozygous null mutants of *apa-2, dpy-23*, or *aps-2* [74, 75, 77]. Strains containing strong loss of function in *apb-1*, however, are embryonic lethal because of the role of APB-1/β1/2 in both AP2 and AP1 complexes, the latter of which function in clathrin-mediated trafficking from the trans-Golgi network and from recycling endosomes [63, 77, 78].

To determine if depletion of the other AP2 subunits could suppress *nekl* molting defects, we carried out RNAi in *nekl-2; nekl-3* animals. RNAi of *apa-2* (α) or *aps-2* (σ) allowed ∼ 40% of *nekl-2*; nekl-3 animals to reach adulthood (Fig 1G). Furthermore, a CRISPR-generated null allele of *apa-2* (*fd280)* led to a slightly higher proportion of *nekl-2; nekl-3* animals reaching the adult stage (Fig 1H, S1 Table). Attempts to suppress *nekl-2; nekl-3* arrest by *apb-1(RNAi)* were unsuccessful due to early embryonic lethality, as expected. Our findings indicate that suppression of *nekl-2; nekl-3* molting defects are not specific to individual AP2 subunits or hemicomplexes, although loss of the µ/β hemicomplex may provide a higher level of suppression than loss of α/σ (Fig 1F,H).

We next determined if suppression of *nekl-2; nekl-3* double mutants by inhibition of AP2 was specific to either the NEKL-2 or NEKL-3 pathways. For these tests, we first carried out *aps-2(RNAi)* in *nekl–mlt* backgrounds containing moderate-to-strong loss-of-function alleles (Fig 1I,J) that exhibit 100% larval arrest as single mutants [20, 73]. Downregulation of *aps-2* led to significant suppression of larval lethality in moderate-to-strong loss-of-function alleles of *nekl-2, nekl-3,* and *mlt-4* (Fig 1I), with weakest suppression observed for *fd90*, a relatively strong *nekl-2* allele. In contrast, no suppression was observed in strains containing a null allele of *nekl-3(gk506)* (Fig I).

Failure to observe suppression of *nekl–mlt* null alleles using RNAi to knockdown AP2 may be due to incomplete inactivation of the targeted subunit. We therefore determined if null alleles in *apa-2*/α or *dpy-23/*µ could suppress larval arrest in null nekl–mlt backgrounds. In the case of *apa-2*, we failed to observe any suppression in null *nekl-2, nekl-3*, and *mlt-3* backgrounds (*n* > 1000 for each strain), indicating that complete loss of the α/σ hemicomplex is not sufficient to overcome the requirement for NEKL-2 and NEKL-3. Attempts to score suppression of *nekl–mlt* null alleles with *dpy-23(e840*), however, were not successful because the generated compound mutants were very sick and slow growing.

### Loss of FCHO-1, an activator of AP2, suppresses *nekl–mlt* defects

Among *nekl-2; nekl-3* animals containing the *fd131* suppressor mutation, 87% progressed to adulthood (Fig 2A,E). Prior genetic characterization of *fd131* indicated the causal mutation to be recessive and autosomal [72]. We determined the *fd131* causal mutation to be a C-to-T transition in the 13^th^ exon of *fcho-1*, which introduces a premature stop codon following amino acid (aa) 903 (Q904Stop; Fig 2C,D, S1 Table). This mutation is predicted to truncate the 963-aa FCHO-1 protein within the conserved Mu-Homology domain (Fig 2C,D). Consistent with this, CRISPR-generated truncations within the Mu-Homology domain (*fd211* and *fd212)* led to a high percentage of viable adults in the *nekl-2; nekl-3* background, as did a null deletion allele of *fcho-1* (*ox477;* Fig 2C–E). In addition, partial knockdown of *fcho-1* by RNAi resulted in ∼ 40% of *nekl-2; nekl-3* mutants reaching adulthood (Fig 2F).

**Fig 2.**
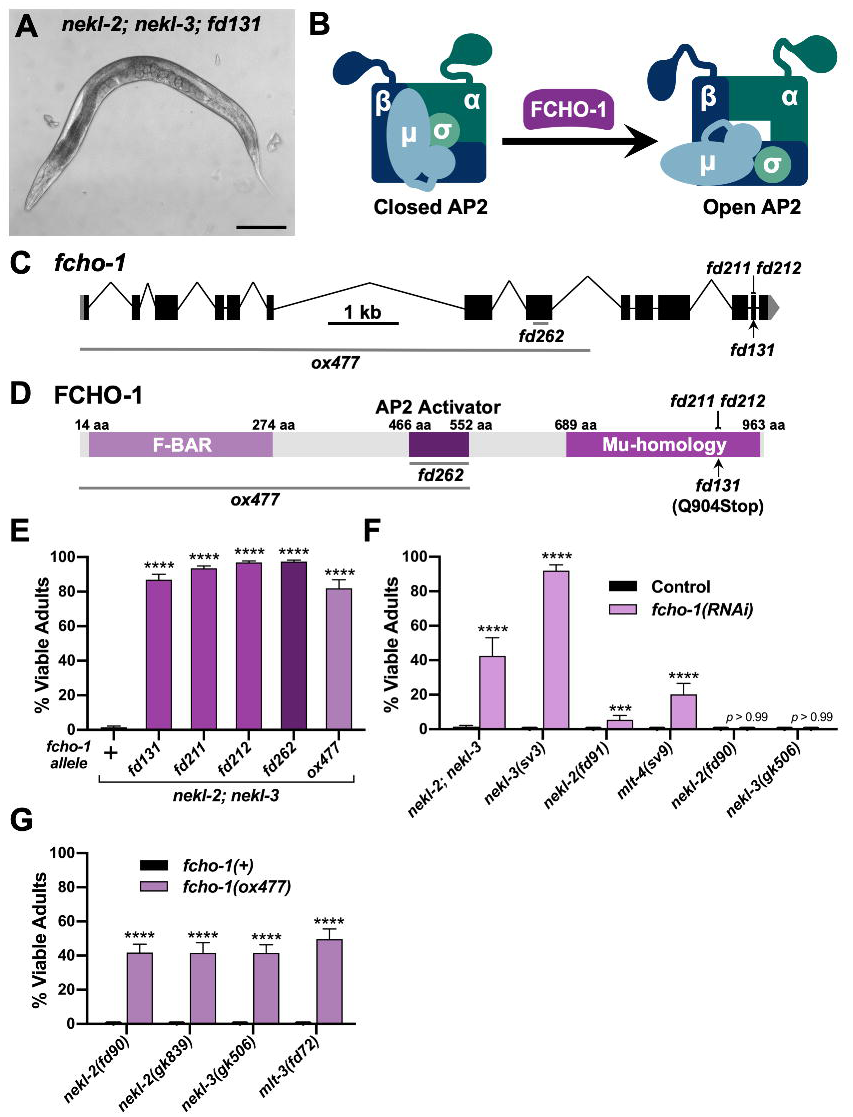
Loss of *fcho-1* activity suppresses *nekl–mlt* molting defects. (A) DIC image of *nekl-2(fd81); nekl-3(gk894345)* adult worm containing the suppressor mutation *fd131*. Bar size in A = 100 µm. (B) Model of AP2 allosteric regulation by FCHO-1. (C) Gene structure diagram of *fcho-1* including the locations of mutations. The point mutation *fd131* is indicated by the arrow, indels (*fd211, fd212*) by the line ending in a small bracket, and deletions (*ox477, fd262*) by horizontal gray lines. (D) Protein domain diagram of FCHO-1 with corresponding allelic changes. The amino acid (aa) locations of the three domains are indicated. (E—G) Bar plots showing suppression of molting defects in *nekl–mlt* mutants by reduction of FCHO-1 activity. (E) Assays were carried out in *nekl-2*(*fd81*); *nekl-3*(*gk894345*) double mutants using the indicated *fcho-1* alleles. (F) RNAi was carried out in the indicated backgrounds using injection methods; control indicates non-injected siblings. (G) Assays were carried out in strong/null *nekl–mlt* backgrounds using the null *fcho-1(ox477)* allele. (E–G) Error bars indicate 95% confidence intervals. p-Values were determined using Fischer’s exact test where proportions were compared to the wild-type allele (E,G) or to the RNAi control (F); \*\*\*\**p* < 0.0001, \*\*\**p* < 0.001. Raw data are available in S1 File.

FCHO-1 is a member of the muniscin protein family, members of which have important roles in clathrin-mediated endocytosis and are recruited to nascent pits early during endocytic vesicle formation [55, 60, 79, 80]. Orthologs of FCHO-1 contain three characterized functional domains: an N-terminal F-BAR domain, a C-terminal Mu-homology domain, and an internal AP2 Activator domain (Fig 1D). The F-BAR domain binds to lipid bilayers and aids in membrane curvature, whereas the Mu-homology domain binds to cargo and other endocytic adapter proteins. The AP2 Activator domain has recently been shown to facilitate the allosteric opening/activation of AP2 (Fig 2B), and loss of *fcho-1* activity in *C. elegans* leads to a decrease in the levels of open/active AP2 [67-69].

Similar to AP2 subunits, suppression of *nekl–mlts* by *fcho-1(RNAi*) was not specific to either the NEKL-2 or NEKL-3 pathways (Fig 2F). Also like the AP2 subunits, *fcho-1(RNAi)* failed to suppress strong loss-of-function alleles of *nekl-2* and *nekl-3 (*Fig 2F). Interestingly, unlike AP2 subunits, a null deletion allele of *fcho-1* (*ox477*) was able to suppress strong/null alleles of *nekl-2, nekl-3*, and *mlt-3* (Fig 2G).

The above findings imply that decreasing the amount of open/active AP2, either by reducing gross AP2 levels or by inhibiting its allosteric activation, can bypass the requirement for NEKL– MLT activity. As a further test of this model, we used CRISPR to generate an in-frame *fcho-1* deletion (*fd262*), which is predicted to specifically remove the AP2 activator domain of FCHO-1 without affecting the F-BAR or Mu-Homology domains (Fig 2C,D) [68, 69]. Importantly, *fcho-1(fd262*) strongly suppressed molting defects in *nekl-2; nekl-3* mutants (Fig 2E), consistent with suppression by *fcho-1* occurring through a reduction in AP2 activity.

### Excess open AP2 enhances *nekl–mlt* molting defects

Given that decreasing the amount of open AP2 led to the suppression of *nekl–mlt* defects, we next tested if increasing the levels of open AP2 could enhance molting defects in weak *nekl–mlt* loss-of-function backgrounds. *ncap-1* encodes the *C. elegans* ortholog of the mammalian adaptiN Ear-binding Coat-Associated Proteins (NECAP1 and NECAP2), which function in several different aspects of clathrin-mediated endocytosis [70, 81-84]. Recently, *C. elegans* NCAP-1 was shown to allosterically regulate AP2 to promote the closed/inactive conformation, and loss of *ncap-1* suppresses the slow-growth phenotype of *fcho-1* strains [70]. Thus, NCAP-1 acts in opposition to FCHO-1 to regulate AP2 conformation and activity (Fig 3A) [67].

**Fig 3.**
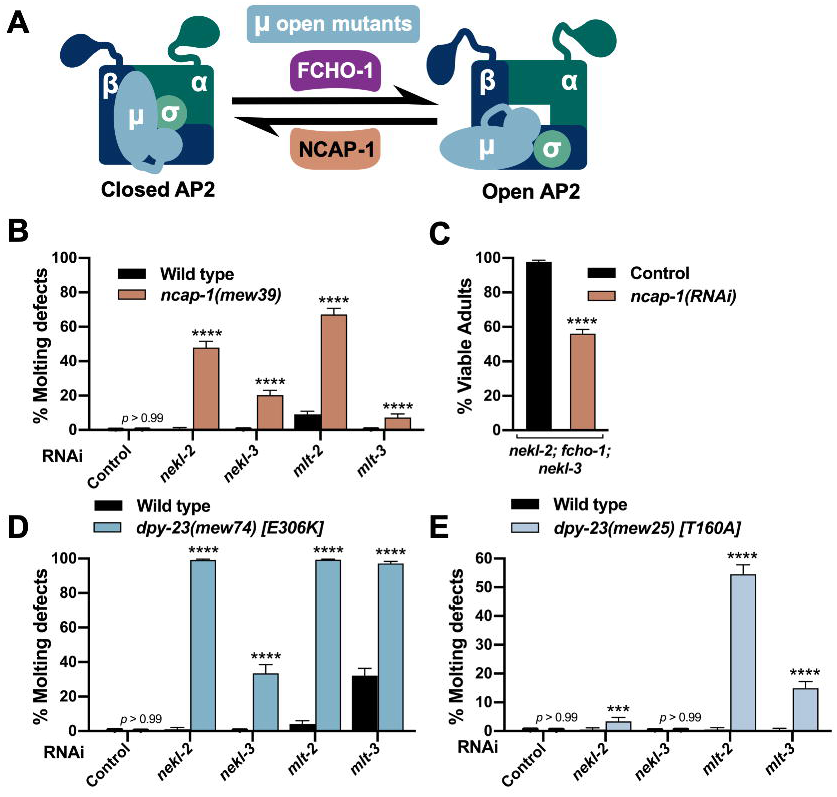
Excess open AP2 enhances nekl*—*mlt molting defects. (A) Model depicting the allosteric regulation of AP2 by FCHO-1 and NCAP-1. DPY-23 (µ subunit) open mutants cause AP2 to remain in the open/active state. (B,D,E) Bar plots showing enhancement of molting defects by mutations in *ncap-1* (B) and *dpy-23* open mutants (D,E). RNAi feeding was carried out for the indicated genes; control RNAi feeding targeted GFP. (C) Bar plot showing partial reversion of suppression in *nekl-2(fd81); fcho-1(fd131); nekl-3(gk894345)* triple mutants after *ncap-1(RNAi)* was carried out using injection methods. Control indicates non-injected siblings. (B–E) Error bars indicate 95% confidence intervals. p-Values were determined using Fischer’s exact test where proportions were compared to the RNAi controls; ****p < 0.0001, ***p < 0.001. Raw data are available in S1 File.

To test for enhancement of molting defects, we carried out RNAi of *nekl–mlts* in wild-type and *ncap-1(mew39*) deletion backgrounds using “weak” RNAi feeding methods, which cause only a partial reduction in *nekl–mlt* activities [20, 21]. Whereas RNAi feeding of *nekl–mlts* in wild type resulted in little or no defective molting, significantly elevated levels of molting arrest were detected in the *ncap-1(mew39*) background (Fig 3B). These findings imply that increased levels of open AP2 exacerbate *nekl–mlt* loss-of-function phenotypes. Correspondingly, RNAi of *ncap-1* mitigated suppression conferred by loss of *fcho-1* in *nekl-2; nekl-3* mutants (Fig 3C). These results are consistent with the reported opposing functions for NCAP-1 and FCHO-1 and support the model that suppression of *nekl–mlts* by fcho-1 and AP2 mutations is due to reduced levels of open AP2.

Because mammalian NECAPs have been suggested to have several distinct functions during endocytosis [81-83, 85], we carried out additional tests to determine if increased open AP2 could enhance *nekl–mlt* defects. For these studies, we made use of several missense mutations in *dpy-23* that shift the balance of AP2 toward the open state (“open mutants”) [68]. As with *ncap-1*, we observed strong enhancement of molting defects after *nekl–mlt* RNAi feeding in *dpy-23* strains containing E306K or T160A substitutions (Fig 3D,E). The somewhat stronger findings observed for the E306K mutation correlates with biochemical assays showing that E306K leads to higher levels of open AP2 than the T160A substitution [68]. Collectively, our findings demonstrate that reducing the level of open AP2 suppresses *nekl–mlt* molting defects whereas increasing the level of open AP2 exacerbates defects.

### Loss of NEKL–MLT activity suppresses AP2-associated defects

Loss of function of non-essential AP2 subunits (µ, α, and σ) and *fcho-1* leads to reduced growth rates and the accumulation of fluid between the cuticle and epidermis [68, 77]. This latter defect visibly manifests in adult-stage worms as bilateral bulges located near the junction of the hyp6 and hyp7 epidermal syncytia in the region of the neck (the “Jowls” phenotype; Fig. 4A). Strikingly, we observed that loss of function in NEKL–MLT activity significantly suppressed the Jowls phenotype of *fcho-1* and AP2 mutants (Fig 4A–F). Specifically, partial suppression of Jowls was observed in the *fcho-1(ox477*) null mutant background, as well as in the presence of null alleles of *apa-2* and *dpy-23,* and in *aps-2(RNAi)* animals (Fig 4C–F). Moreover, suppression was observed in strains with reduced or abolished function in either the NEKL-2 or NEKL-3 pathways (Fig 4C,F). Our observation that simultaneous loss of AP2 and NEKL–MLT activities leads to the mutual suppression of both molting-defective and Jowls phenotypes underscores the tight functional connection between NEKL–MLT and AP2 activities within an intact organism.

**Fig 4.**
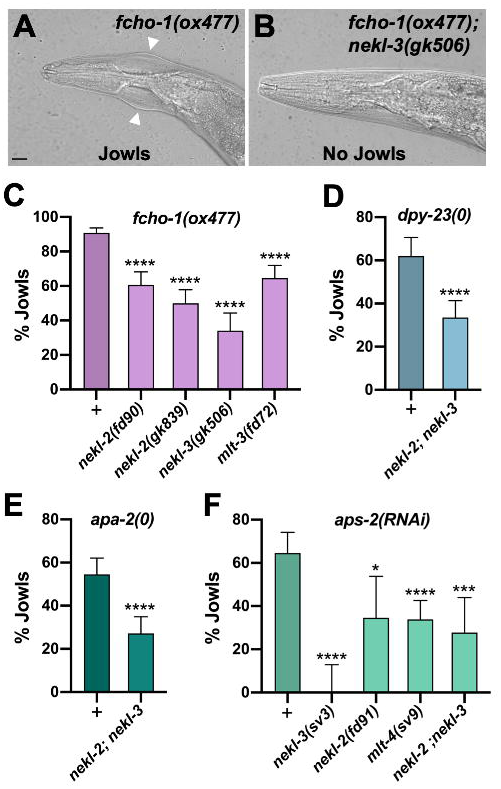
The AP2-associated Jowls phenotype is suppressed by nekl*—*mlt mutants. (A,B) Representative DIC images of *fcho-1(ox477)* (A) and *fcho-1(ox477)*; *nekl-3(gk506)* (B) adults. White arrowheads mark the location of Jowls in *fcho-1(ox477)* mutants. Bar size in A = 5 µm (for A and B). (C–F) The percentage of adult animals exhibiting the Jowls phenotype was assayed in the indicated genetic backgrounds; “+” indicates that no *nekl–mlt* mutations were present. (C) Assays were carried out in strong/null *nekl–mlt* backgrounds using the null *fcho-1(ox477)* allele. (D) Assays were carried out using the *dpy-23* null alleles *fd261* (in the + background), and fd279 (in the *nekl-2*(*fd81*); *nekl-3*(*gk894345*) background). (E) Assays were carried out using the apa-2 null alleles *fd282* (in the + background) and *fd280* (in the *nekl-2*(*fd81*); *nekl-3*(*gk894345*) background). (F) RNAi was carried out in the indicated backgrounds using injection methods. Error bars indicate 95% confidence intervals. p-Values were determined using Fischer’s exact test where proportions were compared to the corresponding wild-type *nekl–mlt* allele (+) (C,D,E) or non-injected controls (F); \*\*\*\**p* < 0.0001, \*\*\**p* < 0.001, \**p* < 0.05. Raw data are available in S1 File.

### Molting-defective *nekl* mutants exhibit changes in epidermal clathrin that are suppressed by inhibition of *fcho-1* and AP2

Given the strong genetic data linking the NEKL–MLTs to AP2, we next examined the role of NEKL–MLTs in clathrin-mediated endocytosis. We previously showed that molting-defective *nekl–mlt* mutants exhibit abnormal localization of a multi-copy clathrin heavy chain reporter [20, 21, 86]. To visualize clathrin at physiological levels, we used CRISPR/Cas9 to generate a clathrin heavy chain reporter construct with an N-terminal GFP tag (GFP::CHC-1), which showed the expected punctate localization pattern in the hyp7 epidermal syncytium (Fig 5A). Consistent with our previous findings, GFP::CHC-1 localization was altered in molting-defective *nekl-2; nekl-3* larvae (Fig 5A,B). Specifically, the average mean intensity of GFP::CHC-1 was increased by 2.2-fold in the apical region of hyp7 in *nekl-2; nekl-3* larvae arrested at the L2/L3 molt relative to wild-type late-stage L2 larvae (Fig 5B). In addition, the percentage of GFP-positive pixels (above a uniformly applied threshold) was increased by 1.3-fold in *nekl-2; nekl-3* molting-defective larvae relative to wild type, which may reflect an increase in the density and/or size of apical GFP::CHC-1 puncta (Fig 5A–C).

**Fig 5.**
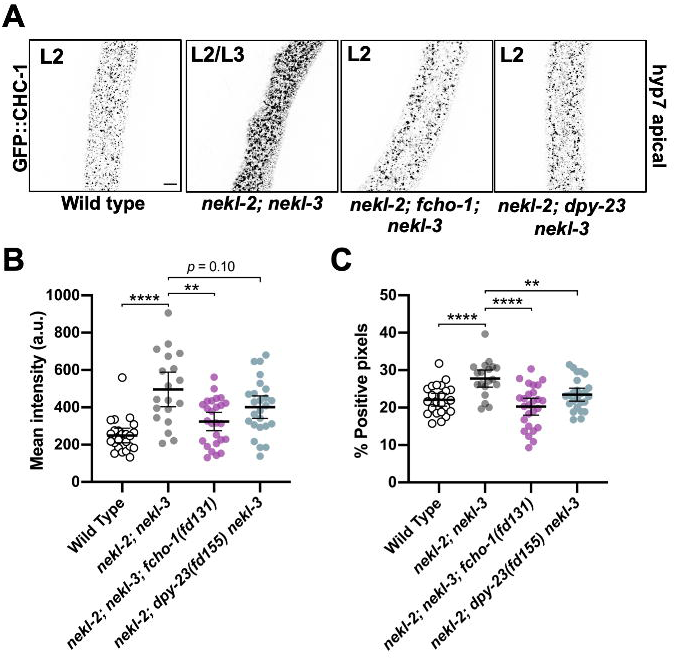
Clathrin defects in *nekl-2*; *nekl-3* mutants are suppressed by reduced AP2 activity. (A) Representative confocal images of L2–L3 larvae expressing CRISPR-tagged GFP::CHC-1 within the apical region of the hyp7 epidermal syncytium. GFP::CHC-1 localization is shown for wild-type, *nekl-2*(*fd81*); *nekl-3*(*gk894345*), *nekl-2*(*fd81*); *fcho-1*(*fd131*); *nekl-3*(*gk894345*), and *nekl-2*(*fd81*); *dpy-23*(fd155) *nekl-3*(*gk894345*) strains. Inverted fluorescence images are shown to aid clarity. Bar size in A = 5 µm (for A–D). Background subtraction was performed using the same parameters for all images; minimum and maximum pixel values were kept consistent for all images. (B,C) For individual larvae of the indicated genotypes, the mean GFP::CHC-1 intensity (B) and the percentage of GFP-positive pixels above threshold (C) were determined in the apical region of hyp7. Both the group mean and 95% confidence interval (error bars) are shown. p-Values for compared means were determined using two-tailed Mann-Whitney tests; ****p < 0.0001, **p < 0.01. Raw data are available in S1 File.

We next determined if clathrin defects in *nekl-2; nekl-3* larvae could be suppressed by a reduction in FCHO-1 and AP2 activities. Relative to *nekl-2; nekl-3* mutants, the average mean intensity of apical GFP::CHC-1 was reduced by 1.5-fold in *nekl-2; fcho-1(fd131); nekl-3* late-stage L2 larvae, and the percentage of positive pixels was reduced by 1.4-fold (Fig 5A–C). A similar trend was observed for *nekl-2; dpy-23(fd155)* nekl-3 animals, which exhibited a 1.2-fold decrease in both the GFP::CHC-1 mean intensity and in the percentage of pixels above threshold relative to *nekl-2; nekl-3* animals, although the observed change in mean intensity was not statistically significant (Fig 5A–C). Together, our findings suggest that reduced AP2 activity mitigates both molting and clathrin-localization defects in *nekl-2; nekl-3* larvae.

### Loss of NEKLs in adults leads to changes in clathrin that are independent of molting defects

Although our above findings suggest that the NEKLs regulate epidermal clathrin, the analysis of trafficking in *nekl–mlt* mutants is problematic because of potential secondary effects caused by the double cuticle and larval arrest phenotype. To circumvent this obstacle, we engineered regulatable NEKL kinases using the auxin-induced degradation system [87, 88]. Importantly, by depleting NEKLs after the final molt, we eliminated the possibility that changes in clathrin localization could be an indirect consequence of defective molting. Proteins tagged with an auxin-inducible degron (AID) are responsive to auxin, which binds to the AID motif leading to ubiquitination by the TIR1–SCF E3-ligase complex and degradation by the proteosome (Fig 6A).

**Fig 6.**
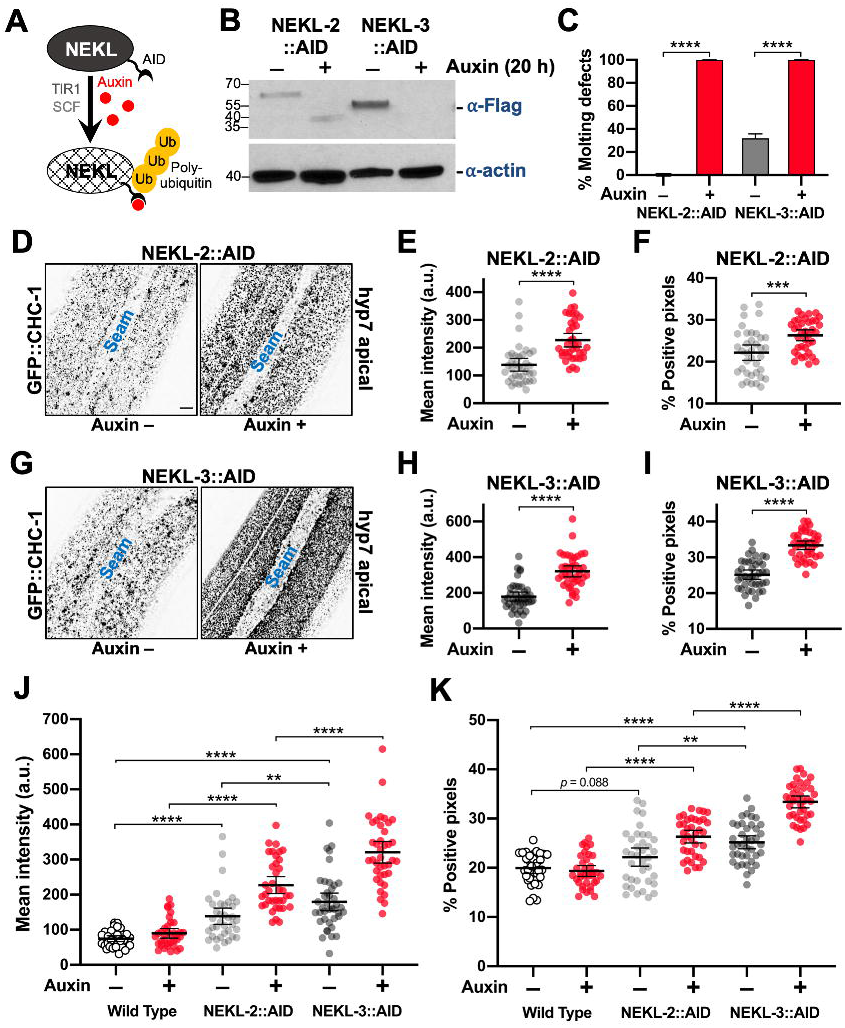
Clathrin localization is altered in NEKL-depleted adults. (A) Model of auxin-inducible degradation of NEKL kinases containing the auxin inducible degron (*NEKL-2*::AID and NEKL-3::AID). The addition of auxin triggers ubiquitination of NEKL::AID proteins followed by proteolysis. (B) Western blot showing complete loss of the full-length *NEKL-2*::AID and NEKL-3::AID proteins after auxin treatment. (C) Bar plot showing the percentage of molting defects among *NEKL-2*::AID and *NEKL-3*::AID strains in the presence and absence of auxin. p-Values were determined using Fischer’s exact test; ****p < 0.0001. (D–K) CRISPR-tagged GFP::CHC-1 localization was analyzed in *NEKL-2*::AID and *NEKL-3*::AID strains in the presence and absence of auxin (20 h) in day-2 adults. (D,G) Representative confocal images of day-2 adults expressing GFP::CHC-1 within the apical region of the hyp7 epidermal syncytium. Background subtraction was performed using the same parameters for all images; minimum and maximum pixel values were kept consistent for all images. Inverted fluorescence was used to aid clarity. Bar size in D = 5 µm (for D and G). (E,F,H,I) For individual adults, the mean GFP::CHC-1 intensities (E,H) and the percentage of GFP-positive pixels above threshold (F,I) were determined for individual adults. (J,K) Summary comparison of data from panels E,F,H,I and S2 Fig (B,C). (E,F,H–K) The group mean and 95% confidence interval (error bars) are shown. p-Values were determined using two-tailed Mann-Whitney tests; \*\*\*\**p* < 0.0001, \*\*\**p* < 0.001, \*\**p* < 0.01. Raw data are available in S1 File.

Exposure of CRISPR-tagged NEKL-2::AID and NEKL-3::AID day-1 adults to auxin led to the complete loss of both full-length tagged proteins within 20 h (Fig 6B). Consistent with NEKL loss of function, 100% of NEKL-2::AID and NEKL-3::AID L1 larvae exposed to auxin from hatching arrested with molting defects (Fig 6C) and displayed abnormal accumulation of apical GFP::CHC-1 (S1 Fig). We note that whereas auxin treatment resulted in the complete disappearance of any detectable NEKL-3::AID protein, we often observed a more rapidly migrating band in samples from auxin-treated NEKL-2::AID worms, suggesting that a partial fragment of NEKL-2::AID may be resistant to further degradation.

To determine the effects of auxin-induced NEKL::AID depletion on clathrin localization in adults, we exposed day-1 adults to auxin for 20 h prior to GFP::CHC-1 localization analysis. In control experiments with wild-type adults, auxin treatment alone did not significantly affect apical hyp7 GFP::CHC-1 localization (Fig 6J,K, S2 Fig). Strikingly, we observed 2.5-fold and 3.6-fold increases in the average mean intensities of apical GFP::CHC-1 in auxin-treated NEKL-2::AID and NEKL-3::AID adults, respectively, relative to auxin-treated wild-type animals (Fig 6D,G,J). Likewise, auxin-treated NEKL-2::AID and NEKL-3::AID adults displayed 1.4-fold and 1.7-fold increases, respectively, in the percentage of pixels above threshold relative to auxin-treated wild-type animals (Fig 6D,G,K). These data demonstrate that loss of NEKLs leads to abnormal apical clathrin localization through a mechanism that is independent of the molting cycle. The stronger effects observed for NEKL-3 could be due to an incomplete loss of NEKL-2 activity after auxin treatment (Fig 6B) or to different requirements for NEKL-2 and NEKL-3 in clathrin-mediated endocytosis.

While conducting these studies, we also observed a clear effect of the AID tag on the activities of NEKL-2 and NEKL-3 even in the absence of auxin. Both mean GFP::CHC-1 levels and the percentage of pixels above threshold were increased in the apical hyp7 region of untreated NEKL-2::AID and NEKL-3::AID adults relative to untreated wild-type controls (Fig 6J,K). The effect of the AID tag was strongest in the NEKL-3::AID strain, consistent with our observation that ∼ 20% of untreated NEKL-3::AID worms displayed molting defects (Fig 6C,J,K). Higher baseline levels in the untreated NEKL::AID strains resulted in somewhat less-dramatic fold changes in comparisons of untreated versus auxin-treated NEKL::AID adults. For example, the average intensity of GFP::CHC-1 increased by 1.6-fold and 1.8-fold in NEKL-2::AID and NEKL-3::AID auxin-treated strains, respectively, compared with age-matched untreated NEKL::AID controls (Fig 6E,H). Correspondingly, the percentage of pixels above threshold was increased by 1.2-fold and 1.3-fold in auxin-treated NEKL-2::AID and NEKL-3::AID adults, respectively, relative to untreated controls (Fig 6,F,I). The observed effects in untreated NEKL::AID strains could be due to the 45-aa AID tag sterically interfering with activities of the NEKL proteins. Alternatively, some auxin-independent activity of the TIR1–E3 ubiquitin-ligase complex could lead to reduced levels of NEKL::AID proteins relative to wild type.

We also assayed GFP::CHC-1 in more medial planes of hyp7, ∼ 1.2 μm from the apical surface, to determine if NEKL depletion led to alterations of clathrin in this region. We observed a modest ∼ 1.2-fold increase in the average mean intensity of GFP::CHC-1 in auxin-treated NEKL-2::AID and NEKL-3::AID adults relative to untreated NEKL::AID worms, although no significant differences were detected when we assayed the percentage of pixels above threshold (S2 Fig). Thus, NEKL depletion has a more pronounced effect on clathrin structures located close to the apical surface of hyp7 than in more medial regions. We also note that wild-type adults showed small but statically significant increases in medial GFP::CHC-1 intensities and pixels above threshold after exposure to auxin (S2 Fig), suggesting that auxin itself may exert a weak effect on GFP::CHC-1 localization. Altogether our findings suggest that loss of NEKLs leads to increased recruitment of clathrin at the apical surface or may extend the lifetime of apical clathrin structures.

### Depletion of NEKLs greatly reduces clathrin exchange on apical membranes of the epidermis

Our results above show overaccumulation of clathrin on apical hyp7 epidermal membranes after depletion of NEKL-2 or NEKL-3. Such an overaccumulation could be due to a higher rate of coated pit formation, a reduced rate of coated pit release from the plasma membrane, or a reduced rate of clathrin-coated vesicle uncoating. One way to differentiate among these possibilities is to measure clathrin exchange between the membrane-associated and cytoplasmic pools via recovery from photobleaching of GFP::CHC-1. If coated pit assembly rates were increased in animals lacking NEKLs we would expect to observe increased recovery rates for GFP::CHC-1 after photobleaching. Conversely, if NEKL depletion leads to a decrease in the disassembly of coated vesicles, as occurs after auxilin depletion, we would expect to observe a decrease in recovery after photobleaching [89]. In addition, certain perturbations that block endocytosis *in vivo* (e.g., exposure to hypertonic sucrose media or intracellular potassium depletion) lead to the genesis of abnormal “clathrin microcages” just below the surface of the plasma membrane, which also block recovery from photobleaching [90]. Moreover, perturbation of the scission enzyme dynamin, which results in failed pinching off of clathrin-coated vesicles, does not affect GFP-clathrin recovery from photobleaching, even though coated pits over-accumulate on the plasma membrane [90]. Therefore, no change in recovery after photobleaching could indicate a role for the NEKLs in vesicle scission.

To determine if the NEKLs affect clathrin dynamics, we carried out fluorescence recovery after photobleaching (FRAP) in NEKL-2::AID and NEKL-3::AID strains ∼ 20 h after exposure to auxin. In the case of wild-type controls, the mobile fraction of GFP::CHC-1 in the apical hyp7 region appeared to be slightly higher in untreated (75%) versus auxin-treated (64%) animals, although this difference was at most marginally significant (p = 0.071; Fig 7A, S3 Fig, S1,2 movies). Strikingly, the GFP::CHC-1 mobile fraction in NEKL-3::AID adults was reduced to only 20% in auxin-treated worms, a more than 3-fold reduction relative to auxin-treated wild-type animals (Fig 7E,F,H, S3 Fig, S3 movie). Similarly, the mobile fraction in NEKL-2::AID strains was just 44% in auxin-treated animals, a 1.5-fold reduction relative to auxin-treated wild type (Fig 7C,F,G, S3 Fig, S4 movie). Thus, loss of either NEKL-2 or NEKL-3 results in a strong reduction in clathrin exchange.

**Fig 7.**
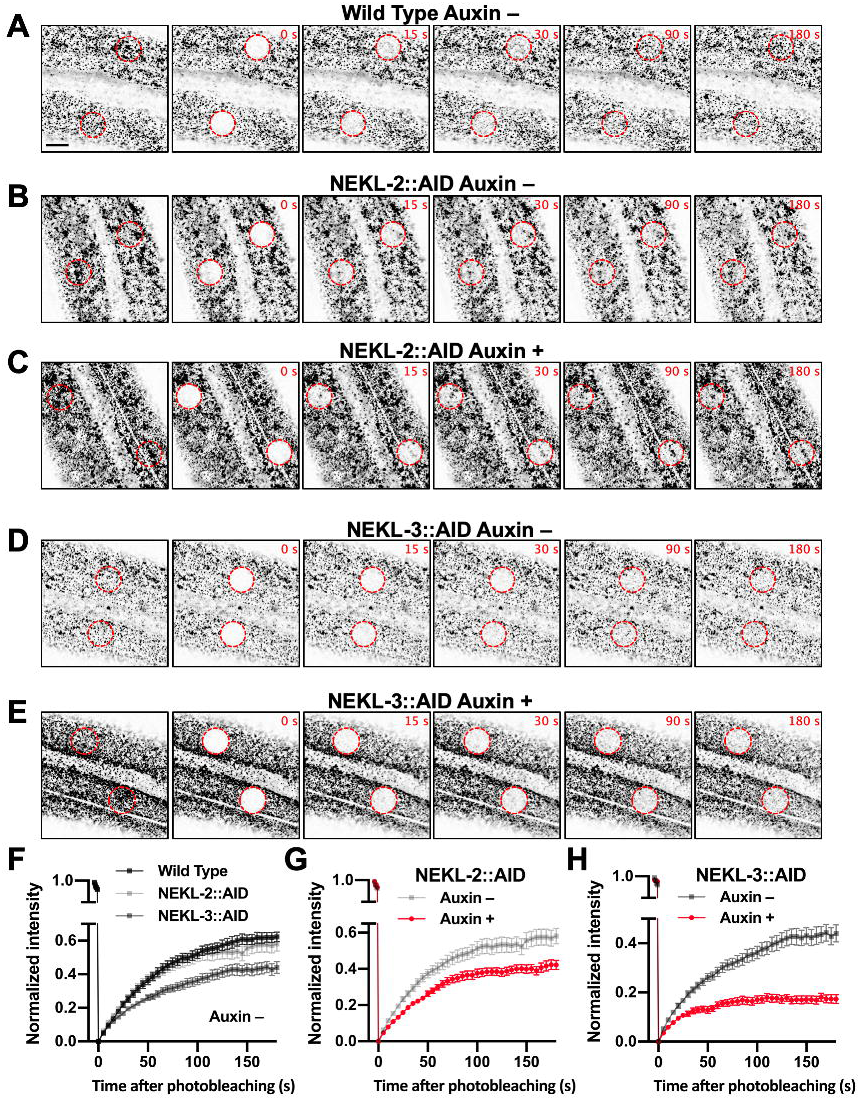
Clathrin dynamics are altered in NEKL-depleted adults. (A—E) Representative time-lapse confocal images of FRAP assays performed using the indicated strains in the presence and absence of auxin (20 h). Images show day-2 adults expressing GFP::CHC-1 within the apical region of the hyp7 epidermal syncytium. Background subtraction was performed using the same parameters for all images; minimum and maximum pixel values were kept consistent for all images. Inverted fluorescence was used to aid clarity. Each panel series contains images at the pre-bleach stage together with images at 0 s, 15 s, 30 s, 90 s, and 180 s after bleaching. Red dashed circles indicate the region of photobleaching. Bar size in A = 5 µm (for A–E). (F–H) Corresponding fluorescence recovery curves of GFP::CHC-1 after photobleaching. Normalized average mean intensities of the photobleached regions were plotted as a function of time using 5-s intervals; error bars denote SEM. (F) Fluorescence recovery curves for wild-type, NEKL-2::AID, and *NEKL-3*::AID adults in the absence of auxin. (G,H) Fluorescence recovery curves for *NEKL-2*::AID (G) and *NEKL-3*::AID (H) strains in the presence and absence of auxin. Raw data are available in S1 File.

We note that similar to what we observed for GFP::CHC-1 mean intensities and pixels above threshold (Fig 6), the AID tag appeared to cause a partial loss of NEKL-2 and NEKL-3 activities, even in the absence of auxin; the mobile fraction in untreated NEKL-2::AID and NEKL-3::AID worms was 61% and 49%, respectively (Fig 7B,D,G,H, S3 Fig, S5,6 movies). In addition, the more robust effects observed for NEKL-3::AID versus NEKL-2::AID adults are consistent with the stronger effects observed for NEKL-3::AID in the GFP::CHC-1 localization assays (Fig 6). Overall, our FRAP data demonstrate that NEKLs strongly affect clathrin exchange in the apical epidermis. Coupled together, our findings indicate that loss of NEKLs leads to the increased stability of clathrin structures located at or near the apical hyp7 surface, a defect consistent with reduced rates of clathrin uncoating and/or the formation of clathrin microcages.

### Clathrin defects in adults are suppressed by reduced activity of AP2

We next determined if clathrin defects associated with adult NEKL::AID loss in adults could be rescued by the *fcho-1* and AP2 suppressor mutations. For these experiments, we examined effects on clathrin in the NEKL-2::AID strain only, as the combination of markers necessary for this analysis in the NEKL-3::AID background led to sickness and slow growth. Consistent with our findings in *nekl-2; nekl-3* larvae (Fig 5), mutations in *fcho-1* and *apa-2* suppressed apical clathrin localization defects in auxin-treated NEKL-2::AID adults (Fig 8A,D). In fact, there were no statistically supported differences between auxin-treated and untreated adults with respect to GFP::CHC-1 mean intensities or the percentage of pixels above threshold in worms containing the *fcho-1* or *apa-2* suppressors (Fig 8B,C,E,F).

**Fig 8.**
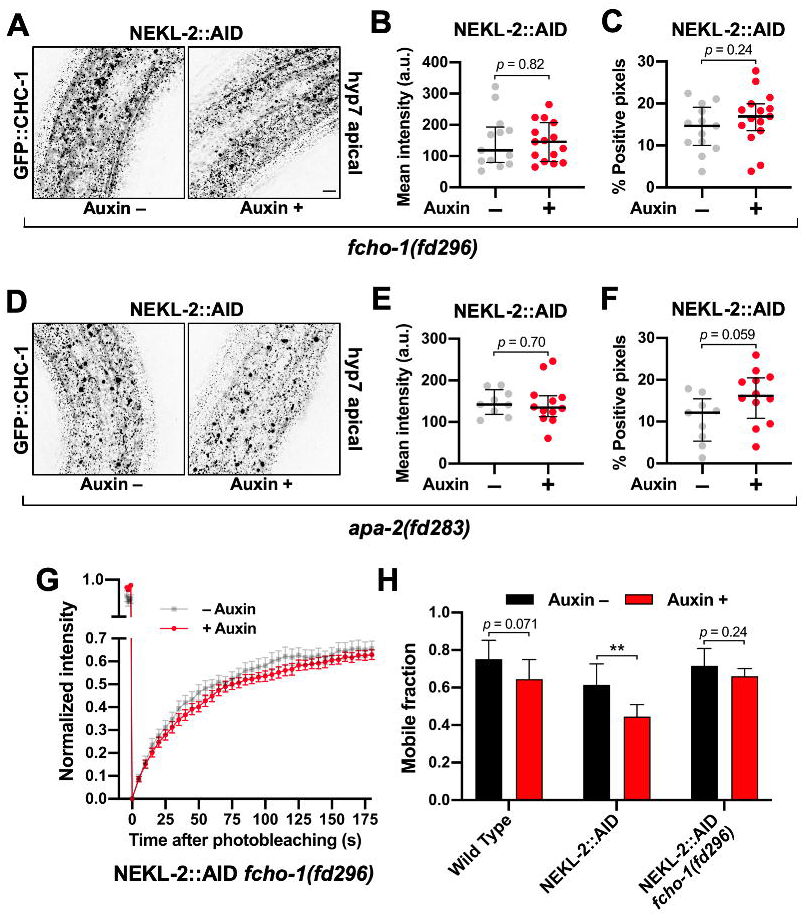
NEKL-associated clathrin defects are rescued by loss of AP2 activity. (A,D) Representative confocal images of day-2 adults expressing GFP::CHC-1 within the apical region of the hyp7 epidermal syncytium. Assays were performed on NEKL-2::AID animals containing null alleles of (A–C) *fcho-1(fd296)* and (D—F) *apa-2(fd283)* in the presence and absence of auxin (20 h). Background subtraction was performed using the same parameters for all images; minimum and maximum pixel values were kept consistent for all images. Inverted fluorescence was used to aid clarity. Bar size in A = 5 µm (for A and D). (B,C,E,F) For individual adults, the mean GFP::CHC-1 intensities (B,E) and the percentage of GFP-positive pixels above threshold (C,F) were determined. (G) Fluorescence recovery curves of NEKL-2::AID *fcho-1(fd296)* day-2 adults in the presence and absence of auxin. Normalized average mean intensities of photobleached regions were plotted as a function of time using 5-s intervals; error bars denote SEM. (H) Bar plot showing the mobile fractions from FRAP analyses of wild-type, NEKL-2::AID, and NEKL-2::AID *fcho-1*(fd296) adults. (B,C,E,F) The group mean and 95% confidence intervals (error bars) are shown. (H) Error bars indicate 95% confidence intervals. p-Values were determined using two-tailed Mann-Whitney tests; **p < 0.01]. Raw data are available in S1 File.

In addition, we assayed GFP::CHC-1 mobility using FRAP in NEKL-2::AID strains containing the *fcho-1(fd296)* suppressor mutation. Untreated day-2 adults exhibited an average mobility of 72%, similar to what we observed for untreated wild type (74%; Fig 7, S3 Fig, Fig 8H). Most notably, auxin treatment had little or no effect on clathrin mobility in NEKL-2::AID adults that contained *fcho-1(fd296*); we observed a 1.1-fold reduction in mobility that was not statistically significant (Fig 8G,H). This is in strong contrast to the 1.5-fold reduction in mobility observed in auxin-treated NEKL-2::AID adults (p < 0.01; Fig 7G, S3 Fig, Fig 8H). Moreover, one-way ANOVA did not indicate statistically significant differences between the GFP::CHC-1 mobilities of wild-type (Auxin +/–), NEKL-2::AID (Auxin –), and NEKL-2::AID *fcho-1(fd296*) (Auxin +/–) adults (p = 0.20). Taken together, our results indicate that AP2 activity can strongly suppress *nekl-* associated clathrin defects in adults independent of larval molting suppression.

### Loss of NEKLs leads to defects in the trafficking of a physiologically relevant membrane cargo

The observed effects on clathrin localization and dynamics following NEKL::AID depletion led us to investigate the role of NEKLs in regulating plasma membrane cargo. LRP-1 is orthologous to human LDL receptor related protein 2 (LRP2), is expressed in hyp7, and is required for normal molting [15]. LRP-1 is thought to bind to and transport low-density lipoproteins into the epidermis from the extracellular space between the epidermis and cuticle. Once internalized, the release and breakdown of LDLs may provide a key source of sterols, which are required for generating hormonal cues necessary for the molting process [4, 13].

Consistent with previous reports, LRP-1::GFP was expressed in a punctate pattern in the apical epidermis of larvae and adults, including in small round puncta that likely represent clathrin-coated pits or vesicles, as well as larger, more irregular accumulations, which are likely to be endosomes (Fig 9A) [15, 91]. Notably, depletion of NEKL-3::AID with auxin led to a dramatic change in the localization pattern of LRP-1::GFP in the apical hyp7 region of adults (Fig 9A,B). Specifically, LRP-1::GFP localization became highly dispersed at the apical membrane after NEKL-3::AID depletion. This effect was reflected by a 2.4-fold increase in the percentage of GFP-positive pixels above threshold in auxin-treated versus untreated NEKL-3::AID adults. Furthermore, no overlap in values between auxin-treated and untreated NEKL-3::AID worms was observed (Fig 9B), indicating that LRP-1::GFP localization provides a particularly strong readout for NEKL defects. Our data suggest that NEKL-3 is required for the maturation of LRP-1−containing clathrin-coated pits and potentially their subsequent delivery from the apical membrane to endosomes.

**Fig 9.**
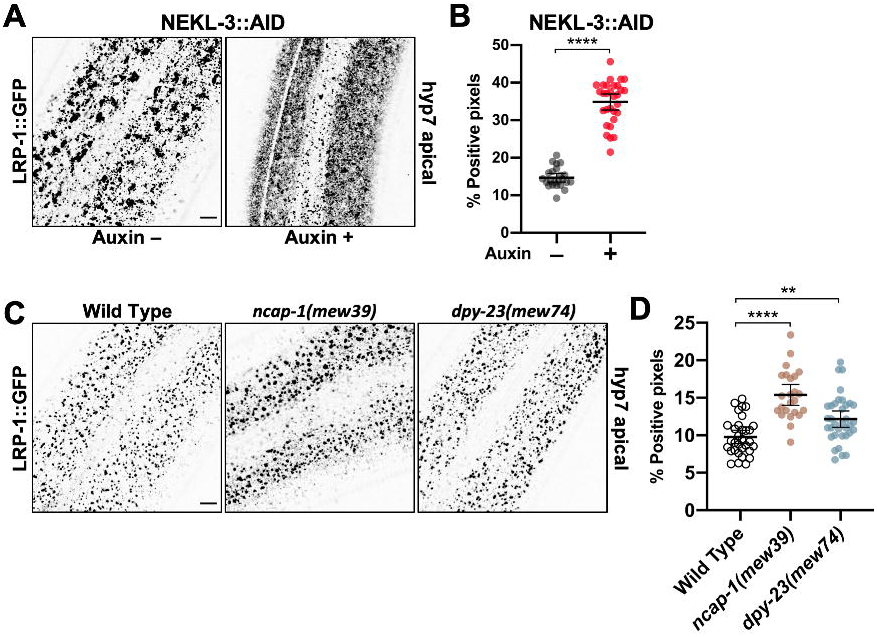
Cargo trafficking is disrupted in NEKL-depleted adults. (A,C) Representative confocal images of day-2 adults expressing LRP-1::GFP in the apical region of the hyp7 epidermal syncytium. Background subtraction was performed using the same parameters for all images; minimum and maximum pixel values were kept consistent for all images. Inverted fluorescence was used to aid clarity. Bar sizes in A,C = 5 µm. (B,D) The percentage of GFP-positive pixels above threshold was determined for individual adults in NEKL-3::AID (+/– auxin 20 h) (B) and wild-type, *ncap-1*(*mew39*), and *dpy-23*(*mew74*) (C) worms. The group mean and 95% confidence interval (error bars) are shown. p-Values were determined using two-tailed Mann-Whitney tests; \*\*\*\**p* < 0.0001, \*\**p* < 0.01. Raw data are available in S1 File.

### NEKLs affect trafficking through a mechanism that is distinct from that of NCAP-1

Several pieces of genetic data suggest that the NEKLs could function through a molecular mechanism that is similar to that of NCAP-1. For example, mutations in *ncap-1* and the *nekls* all suppress the phenotype of AP2 loss-of-function mutants (Fig 4) [70], as do mutations in *dpy-23* that promote the open AP2 conformation [68]. Furthermore, loss of *ncap-1* or *dpy-23* open mutants strongly enhanced molting defects in partial loss-of-function *nekl* backgrounds (Fig 3), which can occur when proteins act in a common pathway or process.

To determine if NCAP-1 and the NEKLs might act through a similar mechanism, we first examined CHC-1::GFP in *ncap-1(mew39*) null mutants. Although mean levels of apical GFP::CHC-1 were increased by 1.2-fold in *ncap-1(mew39*) mutants (Fig 10A,B), these changes were far less pronounced than the ∼ 2- or 3-fold increases observed between auxin-treated wild-type and NEKL::AID adults (Fig 6). Moreover, no statistical difference was detected in the percentage of GFP-positive pixels between wild-type and *ncap-1(mew39*) worms (Fig 10C). Consistent with a lack of strong effects observed for *ncap-1* mutants, we detected no differences in GFP::CHC-1 mean intensity and pixels above threshold between wild type and *dpy-23(mew74)* open mutants (Fig 10A–C). Likewise, we failed to detect statistical differences in clathrin localization in *ncap-1(mew39); dpy-23(mew74)* double mutants, although there appeared to be a trend toward a reduced number of the larger clathrin accumulations (presumed endosomes) in this strain (Fig 10A–C).

**Fig 10.**
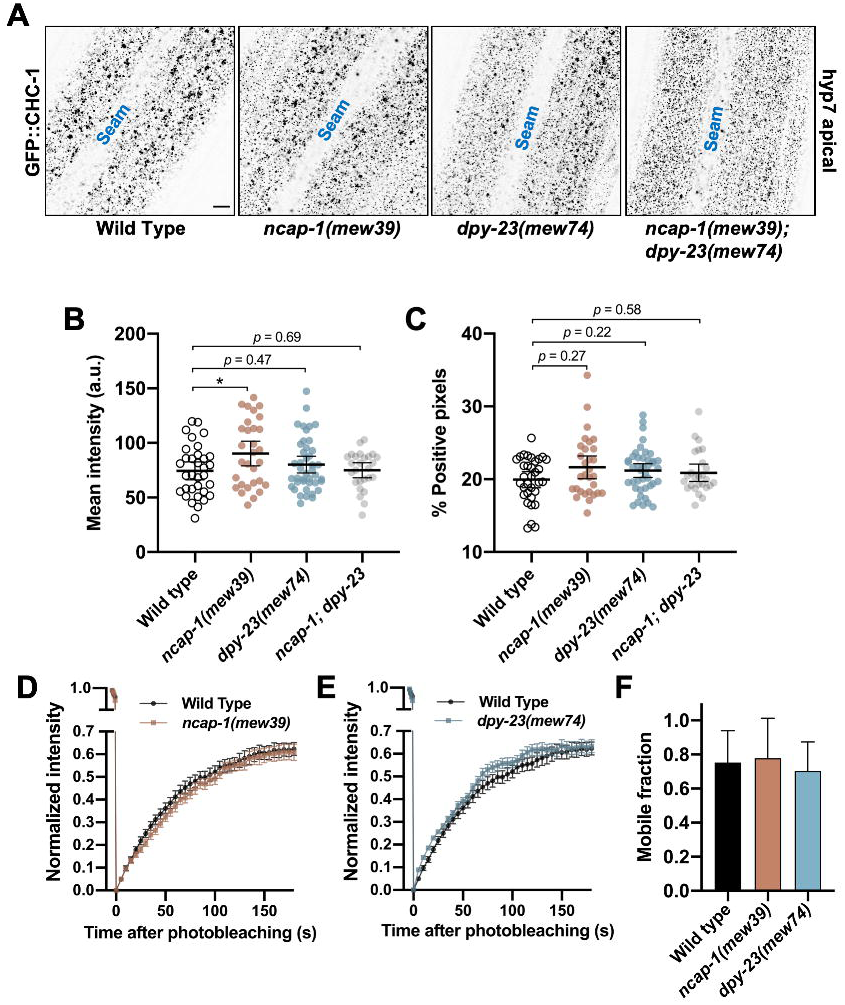
NEKLs and NCAP-1 likely affect trafficking through distinct mechanisms. (A) Representative confocal images of day-2 adults expressing GFP::CHC-1 within the apical region of the hyp7 epidermal syncytium. Background subtraction was performed using the same parameters for all images; minimum and maximum pixel values were kept consistent for all images. Inverted fluorescence was used to aid clarity. Bar size in A = 5 µm. (B,C) For individual adults, the mean GFP::CHC-1 intensity (B) and the percentage of GFP-positive pixels above threshold (C) were determined. The group mean and 95% confidence interval (error bars) are shown. p-Values were determined using two-tailed Mann-Whitney tests; \**p* < 0.05. (D,E) Fluorescence recovery curves of GFP::CHC-1 after photobleaching showing *ncap-1(mew39*) (D) and *dpy-23(mew74)* (E) day-2 adults. Normalized average mean intensities of photobleached regions were plotted as a function of time using −5 s intervals; error bars denote SEM. (F) Bar plot showing the percent mobile fraction from FRAP analyses; error bars indicate 95% confidence intervals. Raw data are available in S1 File.

As an additional test, we examined GFP::CHC-1 recovery after photobleaching in ncap-1(*mew39)* and *dpy-23(mew74)* mutants. Notably, whereas NEKL::AID auxin-treated worms showed a pronounced reduction in the mobile GFP fraction relative to wild type (Fig 7, S3 Fig), we observed no significant differences between wild-type, *ncap-1(mew39)*, and *dpy-23(mew74)* worms (Fig 10D–F). Thus, an increase in the level of open/active AP2 does not appear to result in a marked change in the mobility of clathrin at the apical plasma membrane in hyp7.

We next examined LRP-1::GFP localization in *ncap-1(mew39)* and *dpy-23(mew74)* adults. Although we observed a modest increase in the percentage of pixels above threshold (1.2-to 1.5-fold) in these strains relative to wild type, the effects were again much weaker than what we observed in auxin-treated NEKL-3::AID worms (Fig 9C,D, S4 Fig). These findings strongly suggest that the NEKLs act through a mechanism that is fundamentally different than NCAP-1 and are therefore unlikely to regulate the conformation of AP2 (e.g., to promote the closed state). Nevertheless, our collective data demonstrate that the NEKLs affect a trafficking process that is highly sensitive to the balance between open and closed AP2 conformations.

### The mammalian NEKL-3 orthologs, NEK6 and NEK7, rescue molting and clathrin defects associated with nekl-3 loss

The human NIMA kinase (NEK) family of proteins includes two closely related orthologs of NEKL-3, NEK6 and NEK7, which are ∼ 70% identical and ∼ 85% similar to NEKL-3 and ∼ 80% identical and 90% similar to each other [20]. Given their high degree of conservation, we wanted to determine if the human homologs could rescue molting and clathrin-associated defects *in nekl-3* loss-of-function backgrounds.

We generated transgenes expressing human *NEK6* and *NEK7* cDNAs under the control of the wild-type *nekl-3* promoter. Our constructs included C-terminal fusions of the *NEKs* to an intron-containing GFP cassette as well as cDNA-only constructs that lack the GFP tag. Notably, both *P*_*nekl-3*_::*NEK6::GFP* and *P*_*nekl-3*_::*NEK7::GFP* were able to rescue molting defects in the moderate loss-of-function mutant *nekl-3(sv3)* and in *null nekl-3(gk506)* mutants, although their level of rescue was weaker than what was obtained using a construct expressing NEKL-3 from a 5-kb region derived from the nekl-3 genomic locus (Fig 11A) [20]. In addition, a *NEK6* cDNA-only construct *(P*_*nekl-3*_::*NEK6)* also provided significant rescue in both backgrounds, albeit to a lesser degree than the *NEK6::GFP* fusion. In contrast, a *NEK7* cDNA-only construct *(P*_*nekl-3*_::*NEK7)* failed to rescue molting defects in either background (Fig 11A). The reduced rescuing activity of both the *NEK6* and *NEK7* cDNA-only constructs is likely due to weak expression, as intron-containing markers boost the expression of cDNAs in *C. elegans* and other systems [92, 93]. Consistent with our rescue data for *nekl-3* mutants, *P*_*nekl-3*_::*NEK6*, but not *P*_*nekl-3*_::*NEK7*, was able to partially rescue molting defects following auxin treatment of NEKL-3::AID strains, although rescue was again less robust than for wild-type *nekl-3* (Fig 11B).

**Fig 11.**
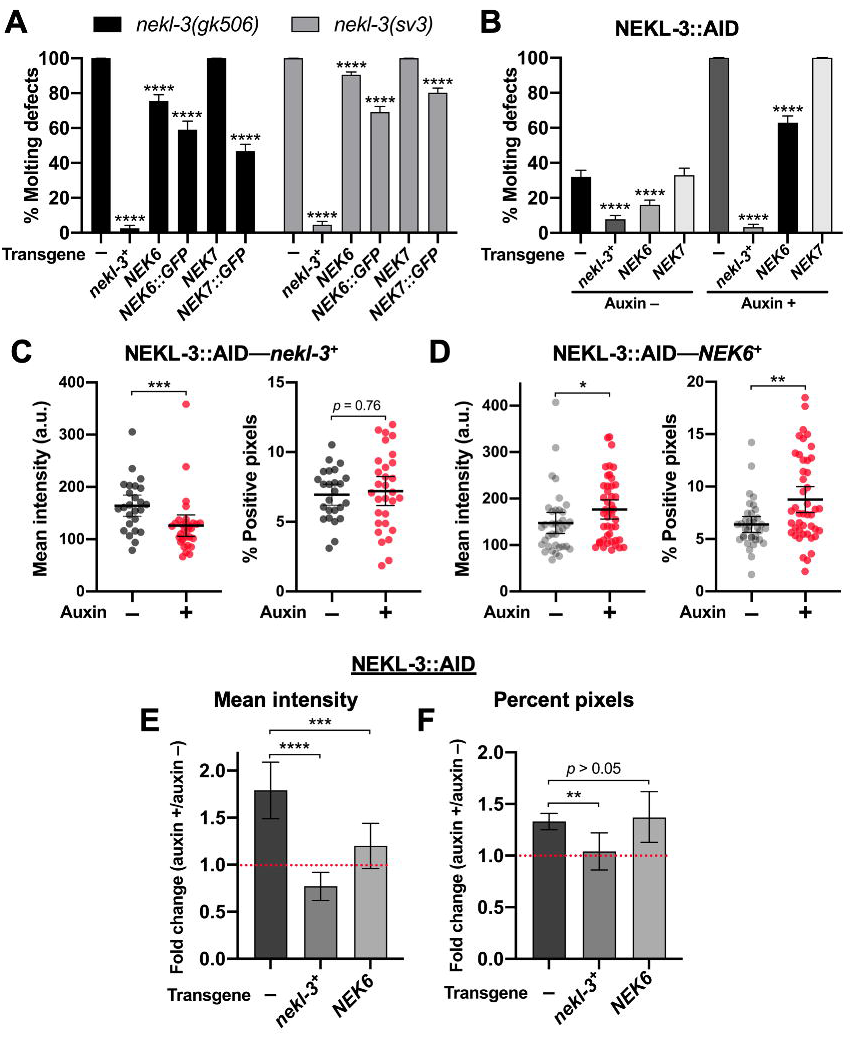
The human orthologs of *nekl-3*, NEK6 and NEK7, rescue molting and trafficking defects. (A,B) Bar plots showing rescue of molting defects in *nekl-3(gk506)* and *nekl-3*(*sv3*) (A) and NEKL-3::AID (B) strains with the indicated transgenes. *NEK6::GFP* and *NEK7::GFP* refer to *P*_*nekl-3*_::*NEK6::GFP* and *P*_*nekl-3*_::*NEK7::GFP*, respectively. *NEK6* and *NEK7* refer to *P*_*nekl-3*_::*NEK6* and *P*_*nekl-3*_::*NEK7*, respectively. p-Values were determined using Fischer’s exact test; \*\*\*\**p* < 0.0001. (C,D) For individual adults, the mean GFP::CHC-1 intensity (left graphs) and the percentage of GFP-positive pixels above threshold (right graphs) were determined for animals expressing wild-type *nekl-3* (C) or *NEK6* (D). The group mean and 95% confidence interval (error bars) are shown. p-Values were determined using two-tailed Mann-Whitney tests; \**p* < 0.05, \*\**p* < 0.01, \*\*\**p* < 0.001. (E,F) Fold-changes (ratios) of auxin-treated (20 h) versus untreated day-2 NEKL-3::AID adults expressing the indicated transgenes were determined for apical hyp7 GFP::CHC-1 mean intensities (E) and the percentage of GFP-positive pixels above threshold (F). Error bars indicate 95% confidence intervals. The dashed red line at 1.0 indicates no change in auxin-treated versus untreated worms. Statistical analyses for ratios were carried out as described in the Materials and Methods; \*\**p* < 0.01, \*\*\**p* < 0.001, \*\*\*\**p* < 0.0001. Raw data are available in S1 File.

We next tested if wild-type *nekl-3* and *P*_*nekl-3*_::*NEK6* could rescue GFP::CHC-1 localization defects in NEKL-3::AID strains. As shown in Fig 6, auxin-induced NEKL-3::AID depletion led to a 1.8-fold increase in mean apical GFP::CHC-1 intensity relative to untreated NEKL-3::AID worms. Furthermore, as expected, expression of the wild-type NEKL-3 protein fully rescued clathrin defects in auxin-treated NEKL-3::AID worms (Fig 11C,E,F). Specifically, no increase was observed in the mean intensity of GFP::CHC-1 or in the percentage of pixels above threshold in treated versus untreated NEKL-3::AID—*nekl-3*^*+*^ worms. Notably, NEKL-3::AID worms expressing NEK6 displayed only a 1.2-fold increase in the mean intensity of GFP:CHC-1 relative to untreated NEKL-3::AID—*NEK6*^*+*^ controls (Fig 11D) and this fold change was significantly lower than NEKL-3::AID worms containing no transgene (Fig 11,E). NEK6 expression did not, however, rescue the percentage of positive pixels above threshold relative to the no-transgene control (Fig 11D,F). In contrast, wild-type *nekl-3* expression fully rescued clathrin defects in both assays (Fig 11D). We note that direct comparisons of GFP::CHC-1 localization data in Figs 6 and 10 were not possible because of differences in marker composition and thresholding procedures. Nevertheless, our findings demonstrate that both human NEK6 and NEK7 can rescue molting defects in *nekl-3* mutants and that NEK6 can partially rescue clathrin defects in strains with reduced NEKL-3 function. These findings strongly suggest that the trafficking functions demonstrated for NEKL-3 are conserved across species.

## Discussion

### NEKLs regulate clathrin-mediated endocytosis

In this study, we have identified the NEKLs as novel regulators of clathrin-mediated endocytosis. More specifically, depletion of NEKL-2 or NEKL-3 at the adult stage led to a strong increase in the levels of clathrin and to a dramatic decrease in the mobility of clathrin at the apical surface of hyp7. These findings demonstrate that endocytic defects following NEKL depletion occur independently of molting defects and are thus not merely a secondary consequence of defective molting. The physiological relevance of our findings is further bolstered by our approach to studying NEKLs within their native context and within an intact developing organism. Moreover, given the requirement for endocytic trafficking factors in the molting process [4], as well as the strong correlation we observed between trafficking and molting defects, our findings indicate that molting defects in *nekl–mlt* mutants are a consequence of abnormal trafficking.

Additional evidence to support a direct role of the NEKLs in trafficking includes our finding that both molting and trafficking defects associated with reduced NEKL–MLT activity are strongly suppressed by mutations that decrease the levels of open/active AP2. This includes loss-of-function mutations of individual AP2 subunits as well as mutations in an allosteric activator of AP2, FCHO-1. Likewise, mutations that increase the levels of open/active AP2 strongly enhanced *nekl–mlt* defects, including *dpy-23/*µ open mutants and loss of function of NCAP-1, an allosteric inhibitor of AP2. Additionally, defects in clathrin localization and mobility after NEKL-2 depletion in adults were rescued by mutations that decrease the levels of open/active AP2. Finally, depletion of NEKL-3 in adults led to dramatic defects in LRP-1/LRP2 endocytosis, which is required for normal molting and is internalized in multiple systems via clathrin-mediated endocytosis [91, 94]. Taken together, our findings indicate that loss of NEKLs leads to defects in the internalization of cargo that is required for normal molting.

### Determining the role of NEKLs in clathrin-mediated endocytosis

One explanation to account for a number of our genetic and cell biological observations is that the NEKLs could promote the closed state of AP2. However, several observations argue against this model. (1) In contrast to clathrin localization in *nekl* mutants and NEKL-depleted adults, clathrin localization in *ncap-1* and *dpy-23*-open mutants was largely unaffected. (2) Unlike clathrin mobility in NEKL-depleted adults, clathrin mobility in *ncap-1* and *dpy-23*-open mutants was indistinguishable from that of wild type. (3) Defects in LRP-1 localization in *ncap-1* and *dpy-23-*open mutants were much less severe than those observed after depletion of NEKL-3. (4) We failed to observe molting defects in *ncap-1(mew39), dpy-23(mew74)* and *ncap-1(mew39); dpy-23(mew74)* double mutants, indicating that increased levels of open/active AP2 alone are not sufficient to induce molting defects. These observations indicate that the NEKLs act through a mechanism that is distinct from that of NCAP-1. Although it remains possible that the NEKLs may carry out multiple functions in endocytosis, including a role in AP2 regulation, we favor a model whereby the NEKLs control a trafficking step that is highly sensitive to the balance between open and closed AP2 conformations.

Based on our FRAP and localization data of clathrin, we suggest that the NEKLs may promote the uncoating of clathrin from internalized coated vesicles. Disassembly of the clathrin coat following membrane scission is carried out by the conserved uncoating ATPase Hsc70, together with its co-chaperone, auxilin [57, 95]. Auxilin bound to clathrin recruits Hsc70 to clathrin-coated vesicles, and Hsc70 then interacts with the clathrin heavy chain to alter the conformation of the triskelia, leading to coat disassembly. Notably, inhibition of DNJ-25, the C. elegans ortholog of auxilin, leads to increased clathrin accumulation within the hyp7 epidermis, decreased clathrin mobility (in coelomocytes), and molting defects [89], all of which are observed in *nekl–mlt* mutants. These observations are consistent with the NEKLs acting either in parallel or upstream of Hsc70–auxilin to promote uncoating, models that will be tested in future studies.

Importantly, we observed that altering the balance between open and closed AP2 conformations led to strong genetic modulation of the molting and clathrin-associated phenotypes in *nekl–mlt* mutants. How might altering the balance of open/closed AP2 impact the efficiency of clathrin uncoating? It is well established that the conformational opening of AP2 promotes clathrin assembly at membranes and the subsequent formation of coated pits [54, 55, 57, 58]. As such, one appealing hypothesis is that the conformational closing of AP2 may in turn facilitate the uncoating of clathrin from internalized vesicles. Although this kind of reciprocal function for AP2 in both the coating and uncoating of clathrin has not, to our knowledge, been directly demonstrated, it is consistent with much of our genetic and cell biological data.

Notably, auxilin can bind to both clathrin and AP2 [96], providing a possible mechanism by which uncoating could be coupled to the conformation of AP2. In addition, it has been suggested that clathrin uncoating may require the disruption of contacts between AP complexes and clathrin [97] and that clathrin and AP2 uncoating are linked [98]. Furthermore, the levels of PIP2 in membranes affect clathrin uncoating [99-101], despite the absence of direct binding of clathrin to membranes. In contrast, AP2 contains a PIP2-binding site that is available for membrane interactions only in the open conformation [55, 66, 67], suggesting that the conformation of AP2 could affect clathrin uncoating.

A number of early studies using cell culture systems, however, demonstrated that clathrin and AP2 uncoating/exchange can take place largely independently of each other. For example, clathrin release can occur in the absence of AP2 release from membranes [102-105]. In addition, cytosolic AP2 can be exchanged with membrane-bound AP2 in vesicles containing a stabilized clathrin coat [98, 106, 107]. Moreover, we failed to observe reduced recovery of clathrin after photobleaching in *ncap-1* and *dpy-23*-open mutants, suggesting that, in an otherwise wild-type background, shifting the AP2 balance toward the open/active conformation does not detectably alter the kinetics of clathrin exchange. Additional studies will be necessary to elucidate the precise mechanism behind AP2 suppression of *nekl–mlt* phenotypes.

We also note that that the reduced mobility of clathrin observed in NEKL-depleted strains could be due to the formation of structures termed “clathrin microcages”. Microcages are extremely small, sharply curved, unusual polymers of clathrin that lack any membrane [108]. Microcages have been observed in mammalian cells that overexpress an inactive form of Hsc70 [109], when potassium or ATP has been depleted from cells, or when cells are exposed to hypertonic sucrose [90, 108]. Similar to what we observed in NEKL-depleted worms, the fluorescence recovery of clathrin after photobleaching was strongly impaired in cells with abundant microcages [90, 98]. These microcages may in fact sequester cytosolic clathrin, leading to reduced clathrin availability and lower levels of endocytosis. Consistent with this, conditions that induce microcages also lead to the dispersal of LDL receptors on the membrane surface [108], a phenotype we observed for the LDL receptor, LRP-1, in NEKL-depleted strains. Future EM studies will determine the ultrastructure of the exchange-resistant clathrin foci in NEKL-depleted worms.

### Suppression of *nekl–mlt* molting defects by actin regulators

We recently published that *nekl–mlt* molting defects can be suppressed by loss of function in the small GTPase CDC-42/CDC42 or by mutations in one of its known effectors, SID-3/ACK1/2 (Activated CDC42 Kinase) [86]. Given that CDC42 and ACK1 are conserved regulators of actin polymerization, this result suggests that an important function of the NEKLs may involve the regulation of actin. Consistent with this, we observed severe defects in actin organization in molting-defective *nekl–mlt* mutants, which were suppressed to varying degrees by inhibition of CDC-42 and SID-3 [86]. Nevertheless, at the time of these studies we were unable to determine if the mislocalization of actin in *nekl–mlt* mutants was a secondary consequence of defective molting or a primary cause of the molting defect.

Although it is possible that suppression of *nekl–mlts* by the actin regulators is unrelated to suppression by AP2, there is extensive data linking CDC42 and actin to a number of endocytic processes including vesicle maturation and invagination, scission from plasma membrane, and trafficking of vesicles away from the plasma membrane [54, 56, 110-118]. In fact, we observed partial rescue of clathrin mislocalization in *nekl-2; nekl-3* mutants treated with *cdc-42(RNAi)* and in *nekl-2; nekl-3 sid-3* suppressed worms [86]. It does, however, remain to be determined if inhibition of *cdc-42* or *sid-3* can suppress clathrin defects in adult-stage NEKL-depleted worms.

Interestingly, there is some evidence to suggest that Hsc70 and auxilin have functions that are connected to actin. For example, Hsc70 binds to actin capping protein, CapZ, which regulates actin filament length [119]. In addition, human auxilin contains a PTEN/tensin domain that interacts with filamentous actin (F-actin) [120, 121], and it has been suggested that auxilin might tether coated vesicles to actin to facilitate clathrin uncoating [122]. We note, however, that the *C. elegans* auxilin ortholog, DNJ-25, does not contain a PTEN/tensin domain and would thus have to interact with actin by another means. Finally, it is worth pointing out that Hsc70 and actin have similar three-dimensional structures and have been implicated in shared functions [123-125]. Taken together, our findings are consistent with the model that the NEKLs may regulate one or more actin-dependent processes that are critical for endocytosis and that *nekl–mlt* molting defects can be suppressed either through mutations affecting actin dynamics or AP2 activity.

Alternatively, it is possible that the NEKLs carry out largely independent functions with respect to actin regulation and intracellular trafficking and that molting defects in *nekl–mlt* mutants result from a combination of defects in both processes. In support of a separate role for NEKLs in cytoskeletal regulation, a striking rearrangement of F-actin occurs during molting, leading to the formation of parallel semi-circumferential actin bundles located at the apical surface of hyp7 [126]. These bundles have been proposed to be important for the molting process, possibly by patterning the structure of the new cuticle. *nekl–mlt* larvae have reduced levels of F-actin and fail to generate parallel actin bundles, a defect that is partially corrected by inhibition of *cdc-42* or *sid-3* [86]. If indeed abnormal molting in *nekl–mlt* mutants is caused by a combination of trafficking and cytoskeletal defects, it would also follow from our data that suppression of either defect is sufficient to rescue the molting phenotype of mutants. Future use of the NEKL::AID system will be particularly helpful in resolving the connection between actin and trafficking and for determining more precisely where and how the NEKLs are acting in these processes.

### Trafficking functions of NIMA kinase family members are likely conserved across species

Our rescue of *nekl-3* molting defects with constructs expressing human NEK6 and NEK7 strongly indicates that these proteins carry our similar functions in diverse organisms (Fig 11). Moreover, we observed rescue of clathrin mislocalization defects in *nekl-3* mutants with human NEK6 (Fig 11). Although the large majority of papers on NEK6 and NEK7 have focused on functions associated with cell division, a high-throughput analysis of the human kinome by Zerial and colleagues indicated that clathrin-mediated endocytosis is strongly decreased in cells when *NEK6* or *NEK7* is targeted by siRNAs [50]. Likewise this study also identified moderate defects in endocytosis following siRNA treatment of *NEK8* and *NEK9*, the closest human homologs of *nekl-2*. A second large-scale study by the Zerial group also reported abnormal trafficking after knockdown of NEK6 and NEK8 based on multiple parameters [51]. In addition, there is some evidence to suggest that NIMA family members in *A. nidulans* and *S. cerevisiae* carry out functions connected to endocytosis [52]. Lastly, protein association studies of NEK6 and NEK7 have identified factors connected to both trafficking (e.g., AP2A1 and AP2B1) and the cytoskeleton (e.g., CDC42 and actin) [36, 127, 128], although the functional significance of these interactions was not explored.

Given these collective observations, we propose that the control of intracellular trafficking may be an ancient and conserved function for members of the NIMA kinase family. This would represent a role for mammalian NEK kinases that has been largely overlooked but could be relevant to their involvement in diseases including cancer and ciliopathies [23, 31, 34, 37-49]. Future studies in both *C. elegans* and mammalian systems will be important to establish the precise functions of NIMA-related kinases in cellular trafficking and human disease processes.

## Materials and Methods

### Strains and maintenance

*C. elegans* strains were maintained according to standard protocols [129] and were propagated at 22°C. Strains used in this study include N2/Bristol (wild type) [130], RT3607 [*pw27(nekl-2::aid); ieSi57(peft-3::mRuby::tir-1); pw17(gfp::chc-1)], RT3608 [pw29(nekl-3::aid); ieSi57(peft-3::mRuby::tir-1); pw17(gfp::chc-1)], RT3402 [pw17(gfp::chc-1)],* WY1145 *[nekl-2(fd81); nekl-3(gk894345); fdEx286 (pDF153(nekl-3 (+)); pTG96)],* WY1271 *[nekl-2(fd81); dpy-23(fd155) nekl-3(gk894345)], WY1531 [dpy-23(fd261)],* WY1560 *[nekl-2(fd81); dpy-23(fd277) nekl-3(gk894345)],* WY1572 *[nekl-2(fd81); dpy-23(fd279) nekl-3(gk894345)],* WY1575 *[nekl-2(fd81); apa-2(fd280) nekl-3(gk894345)],* WY1576 *[nekl-2(fd81); apa-2(fd281) nekl-3(gk894345)],* WY1577 *[apa-2(fd282)],* WY1578 *[pw27(nekl-2::aid); ieSi57(peft-3::mRuby::tir-1); pw17(gfp::chc-1); apa-2(fd283)],* WY1579 *[pw27(nekl-2::aid); ieSi57(peft-3::mRuby::tir-1); pw17(gfp::chc-1); apa-2(fd284)],* WY1580 *[apa-2(fd285) nekl-3(gk506); mnEx174 (F19H6; pTG96)],* EG6353 *[fcho-1(ox477::unc-119(+); unc-119(ed3))],* WY1209 *[nekl-2(fd81); fcho-1(fd131); nekl-3(gk894345)],* WY1350 *[nekl-2(fd81); fcho-1(fd211); nekl-3(gk894345)],* WY1351 *[nekl-2(fd81); fcho-1(fd212); nekl-3(gk894345)],* WY1534 *[nekl-2(fd81); fcho-1(fd262); nekl-3(gk894345)],* WY1592 [nekl-2(fd81); fcho-1(ox477::unc-119(+))]; nekl-3(gk894345)], WY1538 *[nekl-2(fd90); fcho-1(ox477::unc-119(+))],* WY1569 *[nekl-2(gk839); fcho-1(ox477::unc-119(+)],* WY1570 *[fcho-1(ox477::unc-119(+)); nekl-3(gk506)],* WY1571 *[fcho-1(ox477::unc-119(+)); mlt-3(fd72)],* GUN86 *[ncap-1(mew39)],* WY1474 *[dpy-23(mew74)], WY1480 [dpy-23(mew25)],* WY1165 *[nekl-2(fd91[Y84L,G87A]; fdEx278)]*, WY1155 *[nekl-2(fd90[Y84L,G87A,G88A]; fdEx278)],* SP2734 *[mlt-4(sv9); mnEX173(mlt-4(+); pTG96)],* SP2736 *[nekl-3(sv3); mnEx174(F19H6; pTG96)],* LH373 *[nekl-3(gk506); mnEx174(F19H6; pTG96)],* WY1061 *[nekl-2(gk839); fdEx257],* WY1593 *[pw27(nekl-2::aid); fcho-1(fd296)* ieSi57*(peft-3::mRuby::tir-1); pw17(gfp::chc-1)],* WY1578 *[pw27(nekl-2::aid); ieSi57(peft-3::mRuby::tir-1)*; *pw17(gfp::chc-1); apa-2(283)],* WY1510 *[ncap-1(mew39); pw17(gfp::chc-1)],* WY1509 *[pw17(gfp::chc-1); dpy-23(mew74)],* WY1533 *[ncap-1(mew39); pw17(gfp::chc-1; dpy-23(mew74)],* WY1562 *[eqIs1(lrp-1::gfp); [ieSi57(peft-3::mRuby::tir-1); pw29(nekl-3::aid)],* LH191 *[eqIs1(lrp-1::gfp); rrf-3(pk1426)],* WY1573 *[eqIs1(lrp-1::gfp); ncap-1(mew39)],* WY1573 *[eqIs1(lrp-1::gfp); dpy-23 (mew74)],* WY1596 *[nekl-3(gk506); fdEx337(pDF422(NEK7(+)); pTG96.2); mnEx174(F19H6; pTG96)],* WY1615 *[nekl-3(sv3); fdEx351(pDF422(NEK7(+)); pTG96.2); mnEx174(F19H6; pTG96)],* WY1612 *[nekl-3(gk506); fdEx348(pDF244(NEK7::gfp(+)); pTG96.2)],* WY1609 *[nekl-3(gk506); fdEx345(pDF241(NEK6::gfp(+)); pTG96.2)],* WY1608 *[nekl-3(sv3); fdEx351(pDF421(NEK6(+)); pTG96.2)],* WY1602 *[nekl-3(gk506); fdEx343(pDF421(NEK6(+)); pTG96.2)],* WY1401 *[nekl-3(sv3); fdEx316(pDF241(NEK6::gfp(+)); pTG96.2)],* WY1405 *[nekl-3(sv3); fdEx317(pDF245(NEK7::gfp(+)); pTG96.2)],* WY1598 *[ieSi57(peft-3::mRuby::tir-1); pw17(gfp::chc-1); pw29(nekl-3::aid); fdEx339(pDF422(NEK7(+)); pTG96.2)],* WY1539 *[ieSi57(peft-3::mRuby::tir-1); pw17(gfp::chc-1); pw29(nekl-3::aid); fdEx327(pDF421(NEK6(+)); pTG96.2)],* WY1563*[ieSi57(peft-3::mRuby::tir-1); pw17(gfp::chc-1); pw29(nekl-3::aid); fdEx330(pDF153(nekl-3(+)); pTG96.2)].*

### CRISPR/Cas9

CRISPR/Cas9 ribonucleoproteins in combination with the dpy-10 co-CRISPR method [131-133] was used to generate genomic lesions except for the *pw27(nekl-2::aid)* and *pw29(nekl-3::aid)* alleles, which were created using the self-excising cassette method [134]. *pw27(nekl-2::aid)* and *pw29(nekl-3::aid)* guide RNA and repair template plasmids were created from existing plasmids pDD268 and pDD268, respectively [21], using Gibson assembly to directly replace mNeonGreen sequences with AID sequences. Cas9 enzyme was purchased from University of California, Berkeley, whereas, crRNA, tracrRNA, and the repair templates were brought from GE Healthcare Dharmacon, Inc. Briefly, 7.8 μl Berkeley Cas9 (6.4 µg/µl), 0.75 μl 3 M KCl, 0.75 μl 200 mM HEPES (pH 7.4), 5 μl 0.17 mM tracrRNA, 0.8 μl 0.3 mM *dpy-10* crRNA, 2 μl 0.3 mM target crRNA, and 0.75 μl; of distilled water were mixed and incubated at 3°C for 15 minutes. After incubation, 16 µl of *dpy-10* repair template and 1.6 µl of 10 µM target repair template was added, and the mixture was injected into the gonads of adult worms of the desired strains. See S1 Text for information on the exact sequences of oligos used for all CRISPR/Cas9 studies; sequence information on the obtained CRISPR/Cas9 lesions is indicated in S1 Table.

### Transgenic rescue

Transgenic strains used in these studies were obtained by microinjecting ∼ 100 ng/μl of the plasmid of interest and ∼ 50 ng/μl of pTG96.2 *(sur-5::RFP)* into worm gonads [135].

### RNAi

dsRNAs corresponding to *apa-2, dpy-23, aps-2, ncap-1,* and *fcho-1* were generated using standard methods [136], followed by injection at 0.8–1.0 µg/µl into worm gonads (see S1 Text for oligo sequences). For molting enhancement studies, RNAi feeding was performed using bacterial strains from Geneservice following standard protocols [137]. Worm strains were first grown for one (Fig 3B,D) or two (Fig 3E) generations on *lin-35(RNAi)* plates to increase RNAi susceptibility [138]. Gravid adults were then transferred to experimental RNAi plates and allowed to lay eggs for ∼ 24 h, and F1 progeny were scored for molting defects after an additional ∼ 72 h. Because mlt-3(RNAi) induced a high percentage of molting defects in wild type, *mlt-3(RNAi)* bacteria were diluted 1:10 with the control *gfp(RNAi)* strain.

### Image acquisition

Fluorescence images were acquired using an Olympus IX81 inverted microscope with a Yokogawa spinning-disc confocal head (CSU-X1). Excitation wavelengths were controlled using an acousto-optical tunable filter (ILE4; Spectral Applied Research). MetaMorph 7.7 software (MetaMorph Inc.) was used for image acquisition. z-Stack images were acquired using a 100×, 1.40 N.A. oil objective; FRAP time-lapse images were acquired using a 60×, 1.35 N.A. oil objective. DIC images were acquired using a Nikon Eclipse epifluorescence microscope using 10×, 0.25 N.A. and 40×, 0.75 N.A. objectives. Image acquisition was controlled by Openlab 5.0.2 software (Agilent Inc.). Animals were immobilized using 0.1 M levamisole in M9 buffer.

For FRAP assays, anesthetized worms were analyzed immediately after being placed on slides (<10 min). The apical region of hyp7 was brought into focus, and a circular spot (7.272 µm in diameter) was photobleached using an iLas2 system (BioVision Technologies) with a 56-ms pulse of a 405-nm laser set at 50% power. After photobleaching, GFP::CHC-1 FRAP was detected by imaging every 5 s for a total of 180 s.

### Image analysis

To quantify the mean intensity and the percentage of fluorescence-positive pixels above threshold (Figs 5,6,8−11), background fluorescence was subtracted from z-stack images using Fiji software (NIH) available at https://imagej.net/Fiji/Downloads). For a given z-plane of interest, the polygon selection tool was used to demarcate the region of hyp7 followed by mean intensity measurement. The percentage of fluorescence-positive pixels for the region of interest was determined after thresholding, and the same thresholding algorithm was used for strain comparisons.

FRAP time-lapse images were aligned to correct for any movement of the worms by using the “Rigid Body” Transformation method in the StackReg plugin (available at https://imagej.net/StackReg). Next, the time-lapse images were background subtracted and analyzed using the custom-written Stowers plugin (available at http://research.stowers.org/imagejplugins), which can be used with the Fiji software. The photobleached region was selected and mean intensities were quantified before photobleaching and after photobleaching for each time point. Fitting of FRAP curves was performed using batch FRAP fit in the Stowers plugin. Fitted curves were then normalized to prebleach mean intensities and then averaged to obtain final recovery curves. Mobile fractions were quantified using the values obtained from the fitted curves. The mobile fraction is determined by the following equation: amplitude/(prefrap-baseline) (see https://research.stowers.org/imagejplugins/ImageJ_tutorial2.html).

### Auxin treatment experiments

Auxin was purchased as indole-3-acetic acid from Alfa Aesar. In these experiments, L4-stage worms were transferred to plates and left to develop into adults (∼ 20 h). A 0.4 M (100×) stock auxin solution was made by dissolving 0.7 g of auxin in 10 mL 100% ethanol. Each plate containing day-1 adults was treated with a mixture of 25 µl of stock auxin solution and 225 µl of distilled water.

### Statistics

Statistical tests were performed as indicated using software from Prism GraphPad. Statistical tests comparing fold change ratios (Fig 11) were carried out as described by Fay and Gerow [139].

## ACKNOWLEDGEMENTS

Some strains were provided by the *Caenorhabditis* Genetics Center (CGC), which is funded by the US National Institutes of Health (NIH) Office of Research Infrastructure Programs (P40 OD010440). We thank Amy Fluet for editing and the G. Hollopeter Laboratory for strains and scientific input. All authors were funded by GM066868 to DSF. In addition, BJ and VL were further supported by INBRE grant P20 GM103432.

## Fig Supplements

**S1 Fig.**
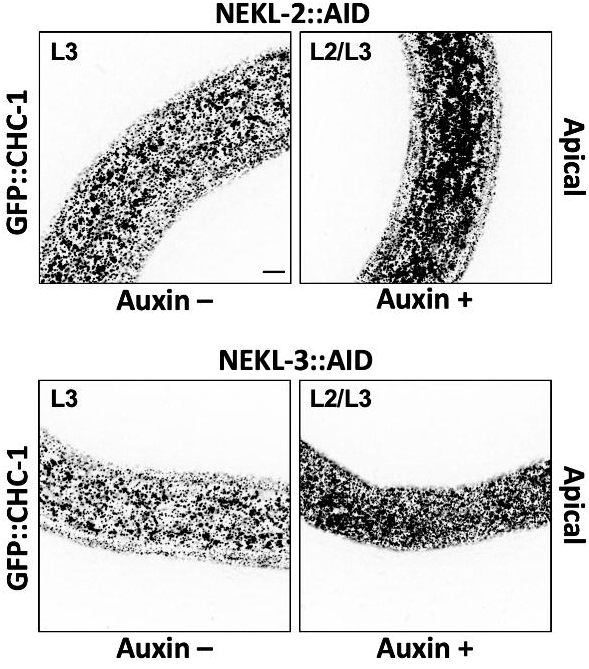
Representative images of untreated (Auxin –) and auxin-treated (Auxin +; 20 h) NEKL-2::AID and *NEKL-3*::AID arrested larvae expressing GFP::CHC-1. Inverted fluorescence images are shown to aid clarity. Background subtraction was performed using the same parameters for all images; minimum and maximum pixel values were kept consistent for all images. Bar in upper left panel = 5 µm (for all panels).

**S2 Fig.**
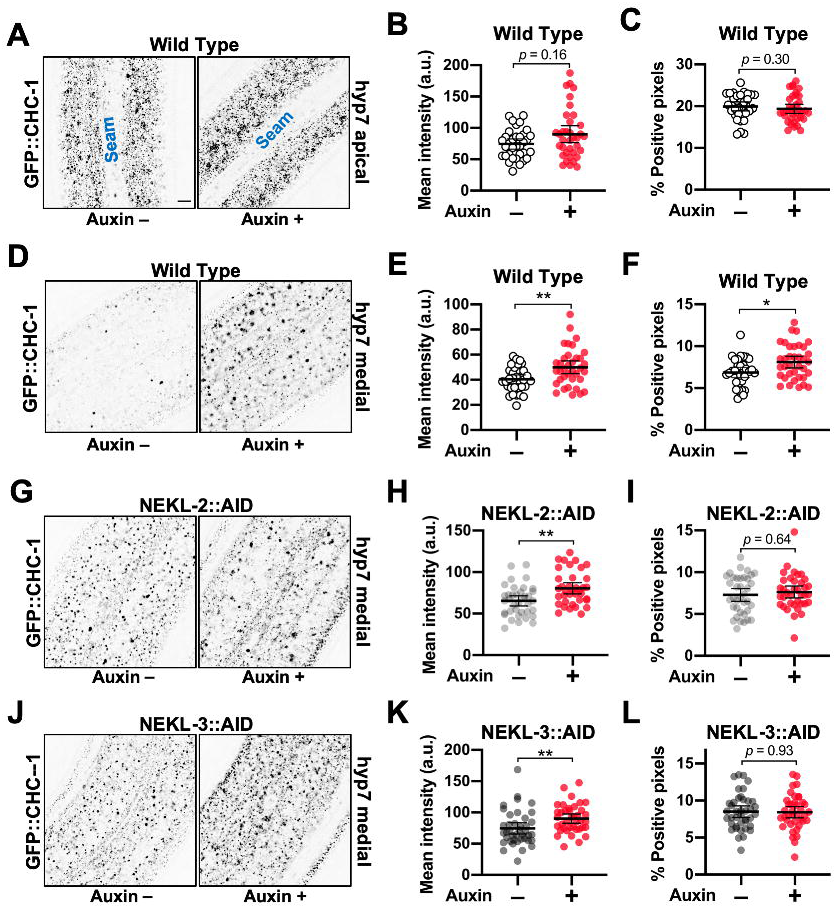
Analysis of GFP::CHC-1 expression in more medial regions of hyp7. (A—F) Representative images of untreated (Auxin –) and auxin-treated (Auxin +; 20 h) wild-type day-2 adults expressing GFP::CHC-1. (G—L) Representative images of similarly treated *NEKL-2*::AID (G—I) and *NEKL-3*::AID (J—L) day-2 adults expressing GFP::CHC-1. Inverted fluorescence was used to aid clarity. Background subtraction was performed using the same parameters for all images; minimum and maximum pixel values were kept consistent for all images. Bar in A = 5 µm (for all panels). Mean GFP::CHC-1 intensities (B,E,H,K) and the percentage of GFP-positive pixels above threshold (C,F,I,L) were determined for day-2 adults. Panels A–C show data for the apical region of hyp7; panels D–L show data for a medial region of hyp7. (B,C,E,F,H,I,K,L) Both the group mean and 95% confidence interval (error bars) are shown. p-Values were determined using two-tailed Mann-Whitney tests; \*\**p* < 0.01, \**p* < 0.05. Raw data are available in S1 File.

**S3 Fig.**
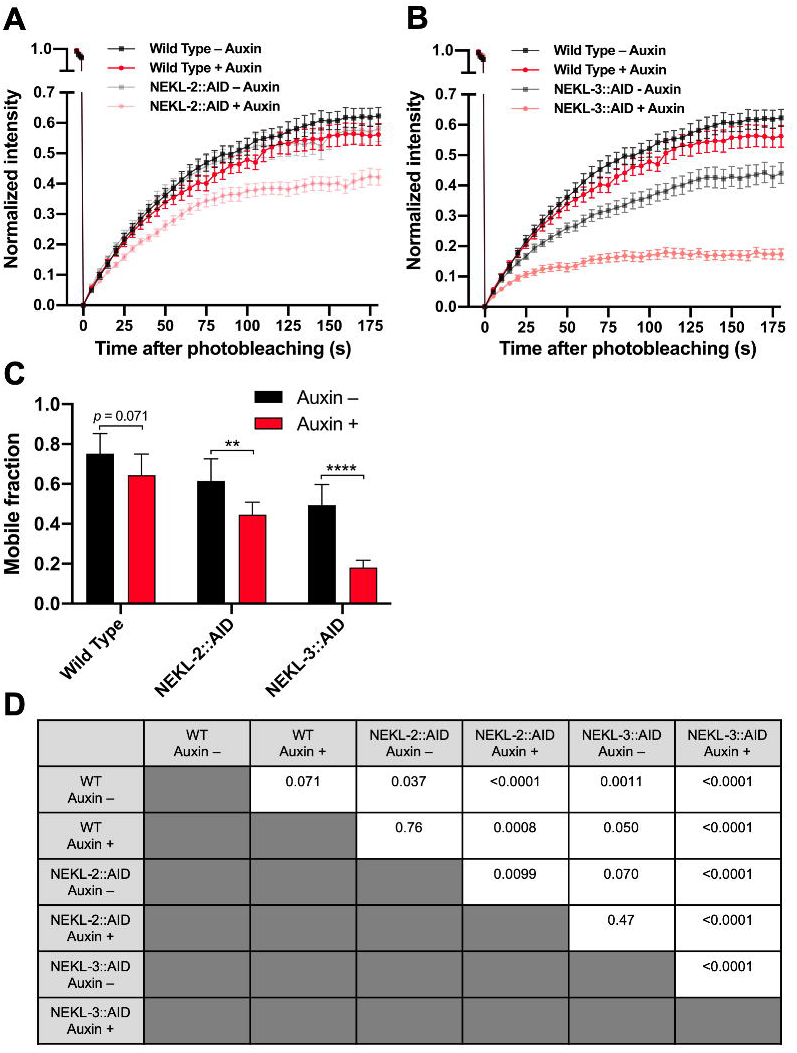
(A,B) Fluorescence recovery curves for wild-type (A,B), NEKL-2::AID (A), and NEKL-3::AID (B) day-2 adults in the presence and absence of auxin (20 h). Analyses were carried in the apical hyp7 region with GFP::CHC-1. Normalized average mean intensities of the photobleached regions were plotted as a function of time using 5-s intervals; error bars denote SEM. (C) Mobile fractions from FRAP data in panels A and B; error bars show 95% confidence intervals. (D) p-Values for all possible comparisons for data in panel C were determined using two-tailed Mann-Whitney tests. Raw data are available in S1 File.

**S4 Fig.**
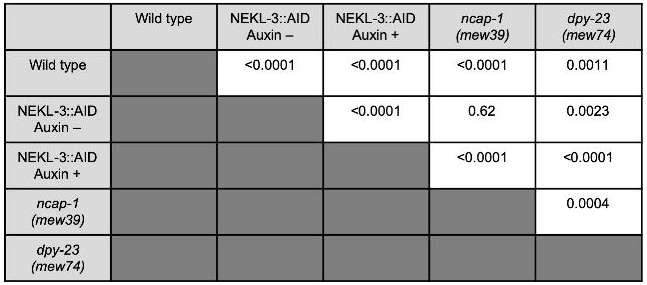
p-Values for all possible comparisons for data in Fig 9 panels B and D (percent positive pixels above threshold) were determined using two-tailed Mann-Whitney tests.

## Supplementary Files

**S1 File.** This excel file contains the raw data used for all quantitative data panels presented in Figs 1–11, including supplementary Figs.

**S1 Text.** This MS Word file contains information describing the generation of all CRISPR alleles used in this study including sgRNAs, repair templates, and sequencing oligos.

**S1 Table.** This MS Word file contains relevant genomic sequencing data of all CRISPR alleles generated in this study.

## Supplementary movies

All Supplementary movies are 200 s long and have been condensed 25× (8 s).

**S1 Movie**. Wild-type (RT3402) day-2 adults; GFP::CHC-1; untreated.

**S2 Movie**. Wild-type (RT3402) day-2 adults, GFP::CHC-1; auxin treated (20 h).

**S3 Movie**. NEKL-3::AID (RT3608) day-2 adults, GFP::CHC-1; auxin treated (20 h).

**S4 Movie**. NEKL-2::AID (RT3607) day-2 adults, GFP::CHC-1; auxin treated (20 h).

**S5 Movie.** NEKL-2::AID (RT3607) day-2 adults; GFP::CHC-1; untreated.

**S6 Movie.** NEKL-3::AID (RT3608) day-2 adults; GFP::CHC-1; untreated.

