## Supplemental Table 1 for "Control of clathrin-mediated endocytosis by NIMA family kinases"

| **CRISPR/Cas9-Modified Regions** | **Gene** |
| --- | --- |
| **Wild Type** (2907) GTAAGCTTGGAAAT_GCTCGATCGGAAGCCGGCGCAGGTCTTCGTGCAATT  ***fd211*** (2907) GTAAGCTTGGAAAT**T**GC A CG AAG**A**C CTTCGTGCAATT | *fcho-1* |
| **Wild Type** (2907) GTAAGCTTGGAAATGCTCGATCGGAAGCCGGCGCAGGTCTTCGTGCAATTT  ***fd212*** (2907) GTAAGCTTGGAAATGCTCGATCGG AGGTCTTCGTGCAATTT | *fcho-1* |
| **Wild Type**(133) GATGAAAAACCGAAGAATCTTCCCGTTGTCGACG­­**-212bp-**AGGAGTACCACTT  ***fd262*** (133) GATGAAAAACCGAAGAATCTTCCCGTTGTCGACG**GTACC**AGGAGTACCACTT | *fcho-1* |
| ***fd296­*** Deletion of undefined length. Mutant exhibits the Jowls phenotype. | *fcho-1* |
| ***fd297*** Deletion of undefined length. Mutant exhibits the Jowls phenotype. | *fcho-1* |
| **Wild Type** (–45)aaagagt­_gattatctcattaaaatacatttcagcaaactcggggATGATTGGT  ***fd261*** (–45) **tt**g gt**g**gatt**g**t tc**g**tt t ttt t­ TGATTGGT | *dpy-23* |
| **Wild Type** (–23) aatacatttcagcaaactcggggATG___AT_T_GGTGGATTGTTCGTTTACA  ***fd279*** (–23) aatacatttcagcaaactcggggATG**TAG**AT**C**T**AA**GTGGATTGTTCGTTTACA | *dpy-23* |
| **Wild Type** (45) ATCAGCGACATCCGAAACTGCAAG AGT A AGGAGGCCGAGCTGAAACGGATC  ***fd280*** (45) ATCAGCGACATCCGAAACTGCAAG**CTT**T**G**A**T**AGGAGGCCGAGCTGAAACGGATC | *apa-2* |
| **Wild Type** (45) ATCAGCGACATCCGAAACTGCAAG AGT A AGGAGGCCGAGCTGAAACGGATC  ***fd281*** (45) ATCAGCGACATCCGAAACTGCAAG**CTT**T**G**A**T**AGGAGGCCGAGCTGAAACGGATC | *apa-2* |
| **Wild Type** (45) ATCAGCGACATCCGAAACTGCAAG AGT A AGGAGGCCGAGCTGAAACGGATC  ***fd282*** (45) ATCAGCGACATCCGAAACTGCAAG**CTT**T**G**A**T**AGGAGGCCGAGCTGAAACGGATC | *apa-2* |
| **Wild Type** (45) ATCAGCGACATCCGAAACTGCAAG AGT A AGGAGGCCGAGCTGAAACGGATC  ***fd283*** (45) ATCAGCGACATCCGAAACTGCAAG**CTT**T**G**A**T**AGGAGGCCGAGCTGAAACGGATC | *apa-2* |
| **Wild Type** (45) ATCAGCGACATCCGAAACTGCAAG AGT A AGGAGGCCGAGCTGAAACGGATC  ***fd284*** (45) ATCAGCGACATCCGAAACTGCAAG**CTT**T**G**A**T**AGGAGGCCGAGCTGAAACGGATC | *apa-2* |
| **Wild Type** (45) ATCAGCGACATCCGAAACTGCAAG AGT A AGGAGGCCGAGCTGAAACGGATC  ***fd285*** (45) ATCAGCGACATCCGAAACTGCAAG**CTT**T**G**A**T**AGGAGGCCGAGCTGAAACGGATC | *apa-2* |

**S1 Table.** Comparison of wild-type sequence and corresponding mutated regions in CRISPR/Cas9 alleles of *fcho-1*(*fd211*, *fd212*, *fd262*, *fd296*, and *fd297*), *dpy-23* (*fd261* and *fd279*), and *apa-2* (*fd280*, *fd281*, *fd282*, *fd283*, *fd284*,and *fd285*). Mutated regions are flanked upstream and downstream by wild-type nucleotides (black font). Numbers indicate distance of the leftmost nucleotide shown from the start of the coding region (ATG) within the cDNA. Stop codons are highlighted in gray. Deleted regions are underlined, insertions are in a blue font, and substitutions are in a red font.
