## Supplementary Text 1 for "Control of clathrin-mediated endocytosis by NIMA family kinases"

**Oligonucleotides used in this study**

For the CRISPR targeting RNAs (crRNAs), target-specific sequences are highlighted in yellow. For repair templates, altered nucleotides are in red, altered restriction endonuclease sites are underlined, inserted nucleotides are highlighted in green, and crRNA target sequences are highlighted in yellow.

***dpy-23* (*fd261*, *fd279*)**

crRNA: 5’- UCAGCAAACUCGGGGAUGAUGUUUUAGAGCUAUGCUGUUUUG-3’

Repair Template:

5’-TTGAAAAGAGTGATTATCTCATTAAAATACATTTCAGCAAACTCGGGGATGTAGATCTAAGTGGA TTGTTCGTTTACAATCACAAAGGAGAAGTGCTCATTTCGAGAATCT-3’

(Creates a BglII site)

Locus Amplification: 5’-TTTGAAGGGCGCTCTATCGG-3’ and 5’-ATGACGTTGACTCGGAAGGC-3’

***fcho-1* (*fd296*, *fd297*)**

crRNA: 5’- ACAGCCGCAAGUCGCCGAGGGUUUUAGAGCUAUGCUGUUUUG-3’

Repair Template:

5’AAGAGGACGTGAATCGAACGAGAAAAGAGCTGAAACAGCCGCAAGTCGCCAGCTGAGTCAATTTGATGCAAACGACGACGACGTGCCTTCAAAAAGCCAAAGAAACG-3’

(Creates a PvuII site)

Locus Amplification: 5’-CCAGGTATCCCACTACATTC-3’ and 5’-CTGGCAGACCAAAATTAGCG-3’

***fcho-1* (*fd262)***

N-terminal crRNA: 5’- CUUCCCGUUGUCGACGACGAGUUUUAGAGCUAUGCUGUUUUG-3’

C-terminal crRNA: 5’- UUGGAAGUAUUACGUUGACG GUUUUAGAGCUAUGCUGUUUUG-3’

Repair Template:

5’-TTGGATGCTCCAATTGATGAAAAACCGAAGAATCTTCCCGTTGTCGACGGTACCAGGAGTACCAC

TTTTGAAAGGGTTCGTTTTTAAGGGGAAAAAATCAATT-3’

(Creates a KpnI site)

Locus Amplification: 5’-AGCATGGACGACACTCACAG-3’ and 5’-GTCTCGCAGACTCATCCAGT-3’

***apa-2* (*fd280*, *fd281*, *fd282*, *fd283*, *fd284*, *fd285*)**

crRNA: 5’-CCGAAACUGCAAGAGUAAGGGUUUUAGAGCUAUGCUGUUUUG-3’

Repair Template:

5’-GATGGCATGCGTGGACTCGCCGTCTTCATCAGCGACATCCGAAACTGCAAGCTTTGATAGGAGGCC

GAGCTGAAACGGATCAACAAAGAATTGGCCAATATTCGATCC-3’

(Creates a HindIII site)

Locus Amplification: 5’-TCAGCGAAACTTTGGAAGCA-3’ and 5’-TGAAGAAAGCAGATTGACGGC-3’

***GFP::chc-1* (*pw17*)**

crRNA: 5’- CUACACACAGUUUUCAAAGAGUUUUAGAGCUAUGCUGUUUUG-3’

Repair Template:

5’-GCAGGTATATAATTTATTGTAAGAAGTTTTTCCATTCTACACACAGTTTTCAAAGATGAGTAAAGGAGAAGAACTTTTCACTGGAGTTGTCCCAATTCTTGTTGAATTAGATGGTGATGTTAATGGGCACAAATTTTCTGTCAGTGGAGAGGGTGAAGGTGATGCAACATACGGAAAACTTACCCTTAAATTTATTTGCACTACTGGAAAACTACCTGTTCCATGGGTAAGTTTAAACATATATATACTAACTAACCCTGATTATTTAAATTTTCAGCCAACACTTGTCACTACTCTCACTTATGGTGTTCAATGCTTCTCGAGATACCCAGATCATATGAAACAGCATGACTTTTTCAAGAGTGCCATGCCCGAAGGTTATGTACAGGAAAGAACTATATTTTTCAAAGATGACGGGAACTACAAGACACGTAAGTTTAAACAGTTCGGTACTAACTAACCATACATATTTAAATTTTCAGGTGCTGAAGTCAAGTTTGAAGGTGATACCCTTGTTAATAGAATCGAGTTAAAAGGTATTGATTTTAAAGAAGATGGAAACATTCTTGGACACAAATTGGAATACAACTATAACTCACACAATGTATACATCATGGCAGACAAACAAAAGAATGGAATCAAAGTTGTAAGTTTAAACATGATTTTACTAACTAACTAATCTGATTTAAATTTTCAGAACTTCAAAATTAGACACAACATTGAAGATGGAAGCGTTCAACTAGCAGACCATTATCAACAAAATACTCCAATTGGCGATGGCCCTGTCCTTTTACCAGACAACCATTACCTGTCCACACAATCTGCCCTTTCGAAAGATCCCAACGAAAAGAGAGACCACATGGTCCTTCTTGAGTTTGTAACAGCTGCTGGGATTACACATGGCATGGACGAACTATACAAATCTGGTGGAGGTGGATCCGCGCTCCCAATCAAATTTCACGAGCACCTGCAGCTTCCGAATGCTGGAATTCGGGTGCCC -3’

**NEKL-2::AID and NEKL-3::AID inserted AID sequence (*pw17* and *pw29*)**

ATGCCTAAAGATCCAGCCAAACCTCCGGCCAAGGCACAAGTTGTGGGATGGCCACCGGTGAGATCATACCGGAAGAACGTGATGGTTTCCTGCCAAAAATCAAGCGGTGGCCCGGAGGCGGCGGCGTTCGTGAAG

**dsRNA Oligos**

For the following dsRNA template primers, target-specific sequences are underlined.

***fcho-1***

5’-TAATACGACTCACTATAGGGAGAGGACGATTGGCGGAACAAAA-3’ and

5’-TAATACGACTCACTATAGGGAGAGTGTCCAGGTTCGAATTTCC-3’

***dpy-23***

5’-TAATACGACTCACTATAGGGAGAACCAAAGAGGAGCAGTCACA-3’ and

5’-TAATACGACTCACTATAGGGAGATCCACACAATGGCATTCTCG-3’

***apa-2***

5’-TAATACGACTCACTATAGGGAGAACATCCGCTGCTTCTCTCAT-3’ and

5’-TAATACGACTCACTATAGGGAGAACGATACCAAACTTCCTCGC-3’

***aps-2***

5’-TAATACGACTCACTATAGGGAGATTCTGATCCAAAACCGTGCC-3’ and

5’-TAATACGACTCACTATAGGGAGAAGGTTATTGTCGGTGATGTC-3’

***ncap-1***

5’-TAATACGACTCACTATAGGGAGAAAAGCATCTCCTCTCTCCTG-3’

5’-TAATACGACTCACTATAGGGAGACATGGGAGATTACGAGAACG-3’
